# Natural selection differences detected in key protein domains between non-pathogenic and pathogenic Feline Coronavirus phenotypes

**DOI:** 10.1101/2023.01.11.523607

**Authors:** Jordan D. Zehr, Sergei L. Kosakovsky Pond, Jean K. Millet, Ximena A. Olarte-Castillo, Alexander G. Lucaci, Stephen D. Shank, Kristina M. Ceres, Annette Choi, Gary R. Whittaker, Laura B. Goodman, Michael J. Stanhope

**Affiliations:** Department of Biology, Temple University, Institute for Genomics and Evolutionary Medicine, Philadelphia, PA 19122, USA; Université Paris-Saclay, INRAE, UVSQ, Virologie et Immunologie Moléculaires, 78352 Jouy-en-Josas, France; Department of Public and Ecosystem Health, College of Veterinary Medicine, Cornell University, Ithaca, NY, 14853, USA; Department of Microbiology & Immunology, College of Veterinary Medicine, Cornell University, Ithaca, NY, 14853, USA; James A. Baker Institute for Animal Health, Cornell University College of Veterinary Medicine, Ithaca, NY, 14853, USA

## Abstract

Feline Coronaviruses (FCoVs) commonly cause mild enteric infections in felines worldwide (termed Feline Enteric Coronavirus [FECV]), with around 12% developing into deadly Feline Infectious Peritonitis (FIP; Feline Infectious Peritonitis Virus [FIPV]). Genomic differences between FECV and FIPV have been reported, yet the putative genotypic basis of the highly pathogenic phenotype remains unclear. Here, we used state-of-the-art molecular evolutionary genetic statistical techniques to identify and compare differences in natural selection pressure between FECV and FIPV sequences, as well as to identify FIPV and FECV specific signals of positive selection. We analyzed full length FCoV protein coding genes thought to contain mutations associated with FIPV (Spike, ORF3abc, and ORF7ab). We identified two sites exhibiting differences in natural selection pressure between FECV and FIPV: one within the S1/S2 furin cleavage site, and the other within the fusion domain of Spike. We also found 15 sites subject to positive selection associated with FIPV within Spike, 11 of which have not previously been suggested as possibly relevant to FIP development. These sites fall within Spike protein subdomains that participate in host cell receptor interaction, immune evasion, tropism shifts, host cellular entry, and viral escape. There were 14 sites (12 novel) within Spike under positive selection associated with the FECV phenotype, almost exclusively within the S1/S2 furin cleavage site and adjacent C domain, along with a signal of relaxed selection in FIPV relative to FECV, suggesting that furin cleavage functionality may not be needed for FIPV. Positive selection inferred in ORF7b was associated with the FECV phenotype, and included 24 positively selected sites, while ORF7b had signals of relaxed selection in FIPV. We found evidence of positive selection in ORF3c in FCoV wide analyses, but no specific association with the FIPV or FECV phenotype. We hypothesize that some combination of mutations in FECV may contribute to FIP development, and that is unlikely to be one singular “switch” mutational event. This work expands our understanding of the complexities of FIP development and provides insights into how evolutionary forces may alter pathogenesis in coronavirus genomes.

## 1. Introduction

Wild and domestic felines worldwide are susceptible to Feline Coronaviruses (FCoVs), with an estimated 12% of infections resulting in deadly Feline Infectious Peritonitis (FIP) (D. Addie et al., 2009). The emergence of mutations within FCoV genomes is thought to be a trigger for FIP development (Pedersen, 2009; Poland et al., 1996; Stoddart & Scott, 1989; Vennema et al., 1998). Significant efforts have been made to compare, often via manual inspection of sequence alignments, genomes obtained from non-pathogenic and pathogenic infections to identify genetic variation that might be associated with FIP development (Brown, 2011). As members of the *Coronaviridae* virus family, FCoVs have some of the largest RNA genomes identified to date (∼29 kb) (Grellet et al., 2022) with some of the highest mutation rates of all evolving systems (Holmes, 2010). Since most viral mutations are expected to have minor phenotypic effects (Frost et al., 2018), identifying those which might impact fitness or pathogenicity requires sensitive statistical tools.

FCoVs belong to the *Alphacoronavirus* genus which also includes coronaviruses (CoVs) that infect dogs (Canine Coronavirus [CCoV]), pigs (Transmissible Gastrointestinal Enteric Coronavirus [TGEV]), and humans (Human Coronavirus 229E [HCoV-229E]) (Li, 2016). More specifically, FCoVs are members of the *Alphacoronavirus 1* species along with CCoV and TGEV (Jaimes et al., 2020). CCoV, TGEV and HCoV-229E can all infect feline cells (Tresnan et al., 1996; Tusell et al., 2007), making felines a potentially important hub for inter-host transmission and virus recombination. There are two unique serotypes that comprise FCoVs, serotype-1 and -2 (FCoV-1 and FCoV-2, respectively). FCoV-2 is thought to be the result of homologous recombination between CCoV serotype 2 and FCoV-1, where FCoV-2 Spike is similar to that of CCoV-2 and the remainder of the FCoV-2 genome to that of FCoV-1 (Herrewegh et al., 1998; Terada et al., 2014). FCoV-1 and -2 each include two biotypes: nonpathogenic Feline Enteric Coronavirus (FECV) predominantly infecting epithelial cells, and pathogenic Feline Infectious Peritonitis Virus (FIPV) robustly infecting macrophages and monocytes (Kipar & Meli, 2014). A tropism shift from epithelial to macrophages/monocytes is a hallmark for FIP development (Kipar & Meli, 2014; Pedersen, 1976; Ward, 1970). The main hypothesis for how FIP develops from an FCoV infection is the “internal mutation” hypothesis, which states that the emergence of virulent, *de novo* mutations from within FECV genomes during infection gives rise to FIPV (H. W. Chang et al., 2011; H.-W. Chang et al., 2010; Herrewegh et al., 1995; Pedersen, 2009; Pedersen et al., 2012; Poland et al., 1996; Stoddart & Scott, 1989; Vennema et al., 1998). The “circulating virulent-avirulent FCoV” hypothesis is less empirically supported, and posits that non-pathogenic and pathogenic strains of FCoV constantly circulate throughout feline populations and FIP results from transmission of the pathogenic biotype (Brown et al., 2009; Healey et al., 2022).

Coronavirus spike proteins are class l virus fusion proteins (Bosch et al., 2003) comprising two subunits, S1 and S2, where receptor recognition is mediated by S1 and membrane fusion by S2 (Li, 2016). The amino (N)-terminal domain (NTD) and carboxy (C)-terminal domain (CTD) of S1 can both act as receptors binding sugar and proteins, respectively (Li, 2016). The main receptor for FCoV-2 is fAPN recognized by the CTD of S1 (Cook et al., 2022; Dye & Siddell, 2007; Tresnan et al., 1996); the main receptor for FCoV-1 remains unknown (Cook et al., 2022; Tekes et al., 2010). It has been demonstrated that the S1 of both serotypes (Spike-1 and Spike-2) can interact with dendritic cell-specific intercellular adhesion molecule grabbing non-integrin (DC-SIGN) acting as a potential co-receptor (Cook et al., 2022; Regan & Whittaker, 2008). Following receptor recognition, but prior to membrane fusion, activation of the Spike protein is required. Activation is often performed by host-cell proteases, e.g., furin (Millet & Whittaker, 2015). FCoV-1 contains two cleavage sites (S1/S2 and S2’), where the S1/S2 site is cleaved by furin (Millet & Whittaker, 2015). FCoV-2 contains only the S2’ site (Millet & Whittaker, 2015). The FCoV-1 Spike S1/S2 furin cleavage site (FCS) is characterized by poly-basic residues S - R - R - S/ A - R - R - S (serine (S), arginine (R), alanine (A)), commonly labeled as P6 - P5 - P4 - P3 - P2 - P1 | P1’, with cleavage occurring between P1 and P1’ (Licitra et al., 2014; Thomas, 2002). Mutations differentiating FECV from FIPV sequences have been identified in this FCS (André et al., 2019; Healey et al., 2022; Licitra et al., 2013, 2014; Millet & Whittaker, 2015; Ouyang et al., 2022). A key feature of class l virus fusion proteins is the proximity of the heptad repeat regions 1 and 2 (HR-1 and HR-2, respectively) to the fusion domain (FD) (Bosch et al., 2003). H.-W. Chang et al., (2012) analyzed FECV-1 and FIPV-1 genomes isolated from infected cats and was the first to report two mutations in the Spike protein – M1058L and S1060A (methionine (M) and serine (S) in FECV and (L) and alanine (A) in FIPV, respectively) that were associated with the shift in virulence. Decaro et al., (2021) reported that FCoVs isolated from 16 of 18 cats diagnosed with FIP contained the M1058L mutation, mirroring what H.-W. Chang et al., (2012) reported. These two mutations fall in the S2 membrane fusion subunit within the connecting region between the FD and HR1. However, the claim that these two mutations are associated with a shift in virulence has been questioned (Barker et al., 2017; Felten et al., 2017; Jähne et al., 2022; Porter et al., 2014), as these mutations have not been found in 100% of FIP cases. Rottier et al., (2005) identified mutations within HR1 and HR2 and suggested that these mutations are responsible for the acquisition of macrophage tropism; a major trigger for FIP development. Several viral accessory proteins, encoded by ORF3abc and ORF7ab, have also been reported to harbor genetic variation associated with the shift in virulence between FECV and FIPV (Brown, 2011), but with discrepancies as to which mutations or deletions within these accessory proteins contribute to the development of the lethal phenotype (Borschensky & Reinacher, 2014; Lutz et al., 2020).

The majority of genetic variation within viral genomes is effectively neutral (Frost et al., 2018). Phenotype-altering mutations, such as those related to drug resistance and immune escape in HIV (Goulder & Walker, 1999; Rambaut et al., 2004), antibody epitopes in Influenza A viruses (Bush et al., 1999), and moderate advantages in infectivity (Hou et al., 2020; Yurkovetskiy et al., 2020), transmissibility (Volz et al., 2021) and convergent evolution of immune evasion (Martin et al., 2021) in SARS-CoV-2 have all been subject to natural selection. Comparative molecular evolutionary analyses of FCoV genomes have the potential to identify phenotype-altering mutations that could be integral to FIP development, thereby pinpointing sites for experimental testing. Xia et al (2020), the only other study we are aware of involving molecular selection analyses of FCoV-1 Spike, identified site 1058 as subject to positive selection in FIPV viral isolates, but did not compare selective regimes of FIPV relative to FECV sequences. Since their publication, statistical methods comparing selection intensities between branch-sets (phenotypes) at sites, as well as gene-wide association of selection with a phenotype (Contrast-FEL (Kosakovsky Pond et al., 2021) and BUSTED-PH (Kosakovsky Pond et al., 2020; Murrell et al., 2015), respectively), have been developed. Furthermore, the use of partial protein coding regions in selection analyses (as was employed in Xia et al (2020)) cannot accurately represent selection acting upon the full-length protein coding region, in turn, limiting the interpretation of results. Therefore, it remains unclear how and where selection is acting differently between both phenotypes.

Herein, we apply comparative statistical techniques to identify sites subject to different selective regimes in FIPV relative to FECV. Further, we identify where selection is associated with the FIPV, FECV, or neither phenotype. We concentrate on full-length protein sequences previously identified to contain the most reported genetic variation between FECV and FIPV sequences – Spike, ORF3abc, and ORF7ab (Brown, 2011). We find two sites evolving differently between FIPV and FECV sequences, as well as 15 sites with evidence of adaptive evolution in FIPV sequences. Eleven of those sites have previously not been reported in literature as associated with the development of lethal disease and warrant subsequent consideration for experimental validation. There also were 38 sites with evidence of adaptive evolution in FECV sequences, 33 of which have not previously been reported as associated with FECV infection.

## 2. Methods

### 2.1 Viral sequence data, genetic recombination, and phylogenetic reconstruction

We queried GenBank (Benson et al., 2018) for all FCoV-1 and -2 protein coding sequences documented to contain the most genetic variation between FECV and FIPV biotypes: Spike, ORF3abc, ORF7ab (Brown, 2011). Overlapping reading frames of ORF3abc and ORF7ab were separated into ORF3a, ORF3b, ORF3c, ORF7a and ORF7b. We manually filtered down the sequence data set based on these criteria:

1. The sequence represents the untruncated, full-length protein coding sequence.
2. The sequence was obtained from a clinical sample collected from a natural infection, and not from an experimental inoculation.
3. The sequence metadata explicitly stated if the sequence was obtained from a clinical FIP diagnosis or not; this information was used to label the sequence as either “FIPV,” or “FECV,” respectively.

All accession numbers of sequences used in our analyses are listed in **Supplementary Table S1**. Sequences that passed the filters were further designated as either serotype-1 or -2, if so annotated. The FCoV-1 and - 2 Spike proteins lack homology across the majority of the S1 domain, and are so distinct that Jaimes et al., 2020 suggested that the two serotypes be thought of as separate viruses. Further, to keep our analyses on full-length protein sequences (*i*.*e*., to refrain from only analyzing the homologous region of Spike), we kept FCoV-1 and -2 Spike analyses separate. We generated codon-aware alignments for each filtered set of protein sequences following the procedure available at the Github repository <u>Codon-MSA</u> (github.com/veg/hyphy-analyses/tree/master/codon-msa). Briefly, in-frame nucleotide sequences were translated, aligned with Multiple Alignment using Fast Fourier Transform (MAFFT) v7.471 (Katoh & Standley, 2013), and then mapped back to corresponding nucleotide sequences. A single copy of identical sequences was retained, resulting in the following number of sequences for each coding alignment: Spike of FCoV-1 (Spike-1): 39, Spike of FCoV-2 (Spike-2): 8, ORF3a: 81, ORF3b: 59, ORF3c: 76, ORF7a: 64, ORF7b: 108.

Evolutionary genetic analyses can be confounded if a single phylogeny is used to analyze a gene alignment, if that alignment has a strong recombination signal, *i*.*e*., where unique topologies are supported by different parts of the gene alignment, typically resulting in higher rates of false positives for selection detection (Kosakovsky Pond et al., 2006). We used the Genetic Algorithm for Recombination Detection (GARD) method (Kosakovsky Pond et al., 2006) to screen alignments for genetic recombination. A maximum likelihood (ML) phylogeny was inferred with RAxML-NG v0.9.0git (Kozlov et al., 2019) under the GTR+Γ nucleotide substitution model for each putatively recombination-free partition (RFP) defined by GARD breakpoints. We used *phylotree*.*js* (Shank et al., 2018) (<u>phylotree.hyphy.org/</u>) to label branches as either “FIPV” or “FECV” in correspondence with metadata. Partitioned protein coding sequence alignments concomitant with the labeled phylogenies served as input for selection analyses and can be downloaded here: <u>data.hyphy.org/web/FCOV/data/</u>.

### 2.2 Detecting differences in natural selection, signals of adaptive, and convergent evolution

We used a variety of codon-based (dN/dS) tests implemented in the HyPhy software package v.2.5.43 (Kosakovsky Pond et al., 2020) to investigate evolutionary hypotheses related to selective pressures differing between FIPV and FECV branches (**Table 1**). All methods were applied to recombinant free partitions (RFP). We used the Contrast-FEL method (Kosakovsky Pond et al., 2021) to identify sites subject to different selective regimes in FIPV relative to FECV sequences. At the gene-wide or RFP-wide level, we compared selection on FIPV relative to FECV labeled branches to identify selection intensity differences with the RELAX method (Wertheim et al., 2015). We modified the BUSTED method (Murrell et al., 2015; Wisotsky et al., 2020) to infer selection on foreground (FG) and background (BG) branches separately (FIPV and FECV branches, respectively), then to statistically associate inferred selection with either the FIPV, or FECV phenotype (<u>BUSTED-PH.bf</u>, (github.com/veg/hyphy-analyses/blob/master/BUSTED-PH/BUSTED-PH.bf).

**Table 1.** Tests applied to detect signals of natural selection.

| **<u>Test</u>** | **<u>Evolutionary unit</u>** | **<u>Method</u>** | **<u>Statistical significance</u>** |
| --- | --- | --- | --- |
| Selective pressure associated with FIPV or FECV or neither | Gene / RFP | **BUSTED-PH** - FIPV vs. FECV | Asymptotic LRT<br>$p \leq 0.05$ |
| Difference in selective pressures between FIPV and FECV | Codon site | **Contrast-FEL** - FIPV vs. FECV | False discovery rate<br>$q \leq 0.20$ |
| Difference in selective pressure intensity between FIPV and FECV | Gene / RFP | **RELAX** - FIPV vs. FECV | Asymptotic LRT<br>$p \leq 0.05$ |
| Episodic diversifying selection | Codon site | **MEME** | Bootstrap (N=500) LRT<br>$p \leq 0.05$ |
| Pervasive diversifying selection | Codon site | **FEL** | Bootstrap (N=500) LRT<br>$p \leq 0.05$ |
| Directional selection | Amino-acid site | **FADE** | Empirical Bayes Factor<br>$EBF \geq 10$ |

If selection was inferred and found to be associated with FIPV, all subsequent site-wise positive selection tests were applied to the FIPV branches, likewise if selection was inferred and found to be associated with FECV branches. If selection was inferred with the BUSTED-PH method, but no significant difference between FIPV or FECV branches was detected, site-wise selection analyses were performed on both FIPV and FECV branches (FCoV-wide). The Mixed Effects Model of Evolution (MEME) and Fixed Effects Likelihood (FEL) methods were used to infer diversifying positive selection (episodic and otherwise, respectively), and FUBAR Approach to Directional Evolution (FADE) (Kosakovsky Pond et al., 2008, 2020) was used to identify directional positive selection (**Table 1**).

We mapped positively selected sites in FCoV-1 Spike to PDB structure 6JX7 (strain UU4 accession number FJ938054) (Yang et al., 2020) using the NGL viewer library to (nglviewer.org/ngl/api/) (Rose & Hildebrand, 2015) in an ObservableHQ Notebook (observablehq.com) (Perkel, 2021), which runs in a web browser. Visualizations can be found here: observablehq.com/@jzehr/fipv1-sites-mapped.

We ran Profile Change with One Change (PCOC) (Rey et al., 2018) (v1.1.0 – github.com/CarineRey/pcoc) on RFPs where BUSTED-PH inferred selection to be associated with the FIPV phenotype, to identify signatures of convergent amino acid evolution across FIPV sequences. We inferred phylogenetic trees from these amino acid alignments using RAxML-NG v0.9.0git (Kozlov et al., 2019) under the PROTGRT model then labeled FIPV branches and used Canine Coronavirus (CCoV) type 1 strain 23/03 (accession number KP849472) to root all trees. Convergent sites were reported with a posterior probability > 0.8.

### 2.3 Protein structural prediction

We generated structural predictions of viral protein ORF3c using AlphaFold2 (Jumper et al., 2021), a deep learning algorithm that leverages multiple sequence alignments and incorporates biological and physical knowledge of protein structures to enable highly accurate predictions of protein structures (Jumper et al., 2021). To reflect the homodimeric nature of a fully intact ORF3c protein, we used AlphaFold-Multimer in ColabFold (Mirdita et al., 2022). Predicted local distance difference test (pLDDT) and predicted aligned error (PAE) were used to quantify confidence in the predicted structure. N-terminal (aa 1-22) and C-terminal (aa 217-236) predicted secondary structure extensions with low pLDDT confidence scores (<50) were not displayed. We used this predicted structure to compare it with the cryo-EM structure of SARS-CoV-2 ORF3a (PDB: 6XDC) and to map the positively selected site located at position 165.

## 3. Results

### 3.1 Genomic Recombination

Coronaviruses (CoVs) are known to extensively recombine (Banner & Lai, 1991; de Klerk et al., 2022; Graham & Baric, 2010; Liao & Lai, 1992; Lytras et al., 2022). Since recombination can confound evolutionary genetic analyses if not properly accounted for (Kosakovsky Pond et al., 2006), we screened each codon-aware alignment for evidence of recombination. Phylogenetic incongruence, a hallmark of recombination, was found in the two Spike serotype -1 and -2 (Spike-1 and Spike-2, respectively) codon-aware alignments resulting in 13, and 8 inferred recombination-free partitions (RFPs), respectively. Breakpoints inferred for each protein can be found in **Supplementary Table S2**. There were no supported breakpoints inferred in ORF3a, ORF3b, ORF3c, ORF7a, and ORF7b.

### 3.2 Natural selection differences between FIPV and FECV phenotypes

We used the Contrast-FEL method (Kosakovsky Pond et al., 2021) to identify differences in natural selection pressures at individual codons between FIPV and FECV sequences. We found two sites subject to detectably different selective pressures in Spike-1 between the two phenotypes (false discovery rate, FDR ≤ 0.2): codon positions 789 and 1046; site 1046 maps to site 1058 first reported by H.-W. Chang et al., (2012) **(Fig. 1A)**. In both cases, a higher nonsynonymous (dN) to synonymous rate (dS) ratio (indicative of stronger positive selection) was detected in FIPV relative to FECV labeled sequences. Site 789 falls within the S1 subunit, in the S1/S2 furin cleavage motif mapping to the P4 position of this poly-basic motif. Site 1058 falls within the connecting region between the fusion domain (FD) and the heptad repeat region 1 (HR1) in the S2 subunit. H.-W. Chang et al., (2012) identified amino acid site 1060 as also associated with the pathogenic shift from FECV to FIPV, however we did not identify measurably different selection at this site between the phenotypes. All sites reported for Spike-1 are ungapped positions in accession FJ938054, strain UU4. All codon and amino acid sites identified and reported herein refer to the ungapped index in the respective reference sequence.

**Figure 1.**
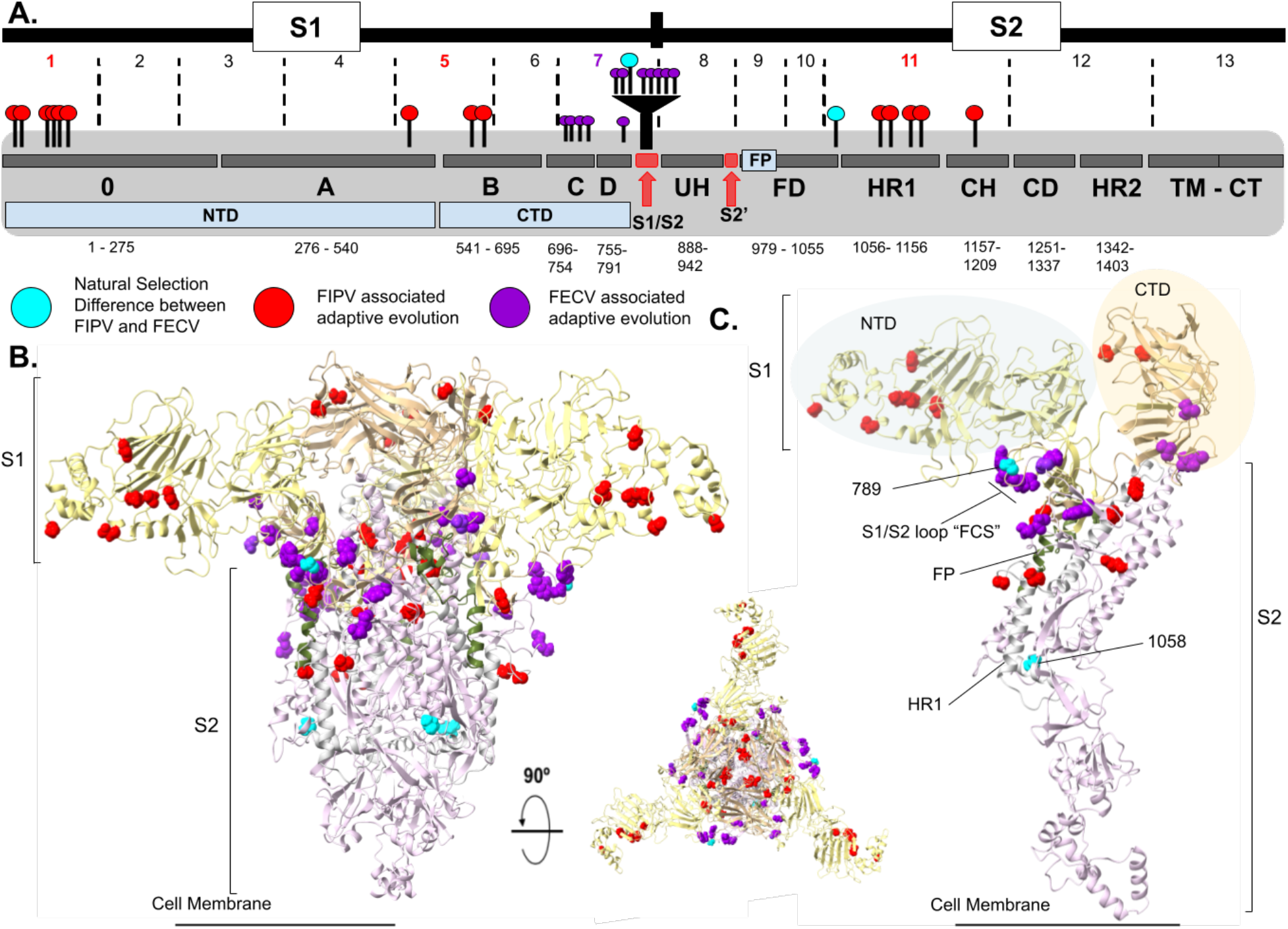
FIPV-1 Spike (Spike-1) domain map and tertiary structure highlighting sites subject to natural selection. Sites are mapped to the protein domain map and PDB structure 6JX7 accession number FJ938054 (Yang et al., 2020). **A)** S1 and S2 subunits of Spike further separated into functional protein subdomains. Dashed vertical black lines delimit numbered RFPs, and are colored based on association of phenotype with inferred selection. The two sites identified by Contrast-FEL (Kosakovsky Pond et al., 2021) to be evolving differently between FIPV and FECV are depicted in cyan. Codon sites subject to adaptive evolution associated with the FIPV phenotype are depicted in red. FECV associated codon sites subject to adaptive evolution are represented in purple. Text labels for each domain: 0-domain; A domain; B domain, receptor binding motif (RBM); C domain; D domain; S1/S2 furin cleavage site (FCS); upstream helix (UH); S2’ cleavage site; fusion domain (FD) with fusion peptide (FP); heptad repeat region 1 (HR1); central helix (CH); connector domain (CD); heptad repeat region 2 (HR2); transmembrane domain (TM); cytoplasmic tail (CT). Amino acid indices are reported for each domain. **B)** Sites mapped to PDB 6JX7 (trimer) to visualize selected sites in 3D space. **C)** The monomer representation. NTD is highlighted in yellow, CTD in gold, FP in green, HR1 in white, and the rest of the S2 subunit in light pink.

The BUSTED-PH (Kosakovsky Pond et al., 2020; Murrell et al., 2015) method was used to infer selection at the gene-wide level and to associate inferred selection with a phenotype (FIPV and FECV) (see Methods for further details). Where gene-wide positive selection was inferred and statistically associated with the FIPV phenotype, 14 sites were identified to be subject to positive selection in Spike-1. These sites were scattered across several functional subdomains, including the 0-domain, B, HR1 and CH, with a particular concentration in the 0-domain (**Fig. 1**). Only a single codon in Spike-2 (site 1404 in accession number X06170 – FIPV strain 79-1146) was judged to be under positive selection and associated with the FIPV phenotype. There were 12 codon sites inferred to be under positive selection associated with the FECV phenotype within Spike-1, almost exclusively within the S1/S2 and C domain **(Fig. 1)**. The two sites subject to positive selection in Spike-2 associated with the FECV phenotype mapped to the Spike-2 RBM (Reguera et al., (2012); **Table 2**). ORF7b selection was associated with FECV, and included a total of 24 sites (**Table 2**). Individual codon sites subject to adaptive evolution in all other partitions where selection signals could not be statistically associated uniquely with one phenotype (i.e. FCoV-wide selection) are reported in **Supplementary Table S3**, and includes: 53, one, seven, three, six, and one sites across the remaining RFPs in Spike-1, Spike-2, ORF3a, ORF3b, ORF3c, and ORF7a respectively.

**Table 2.**
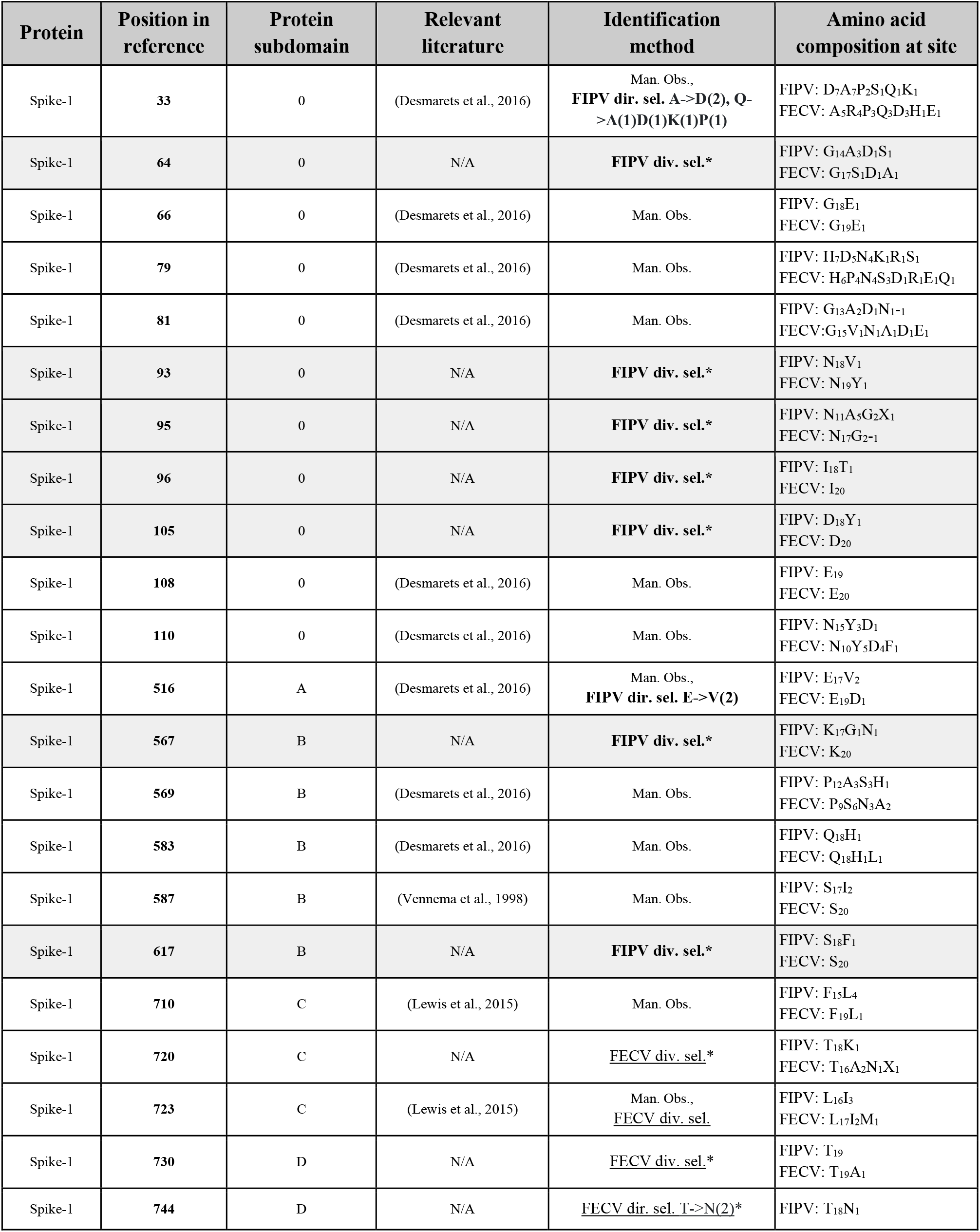
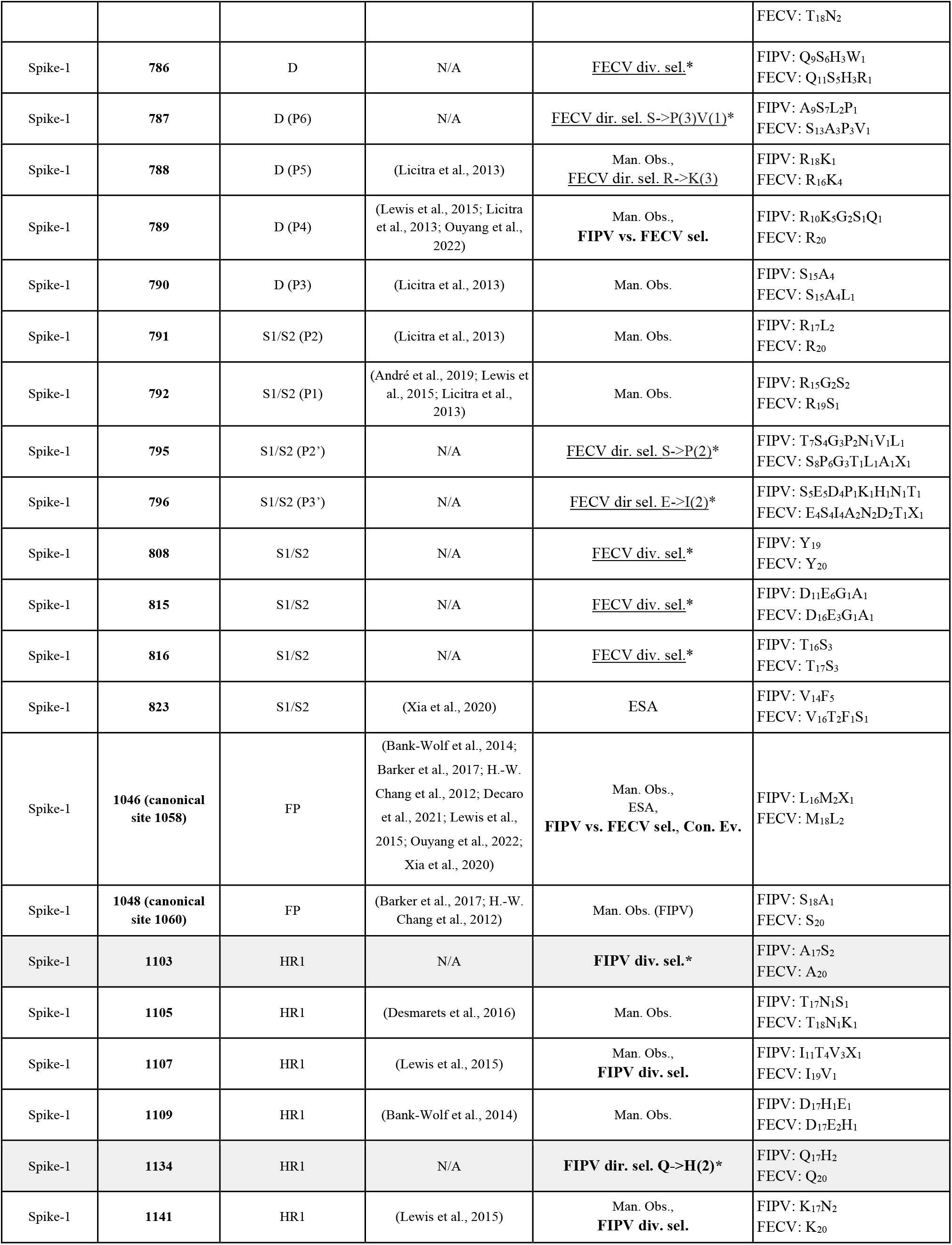
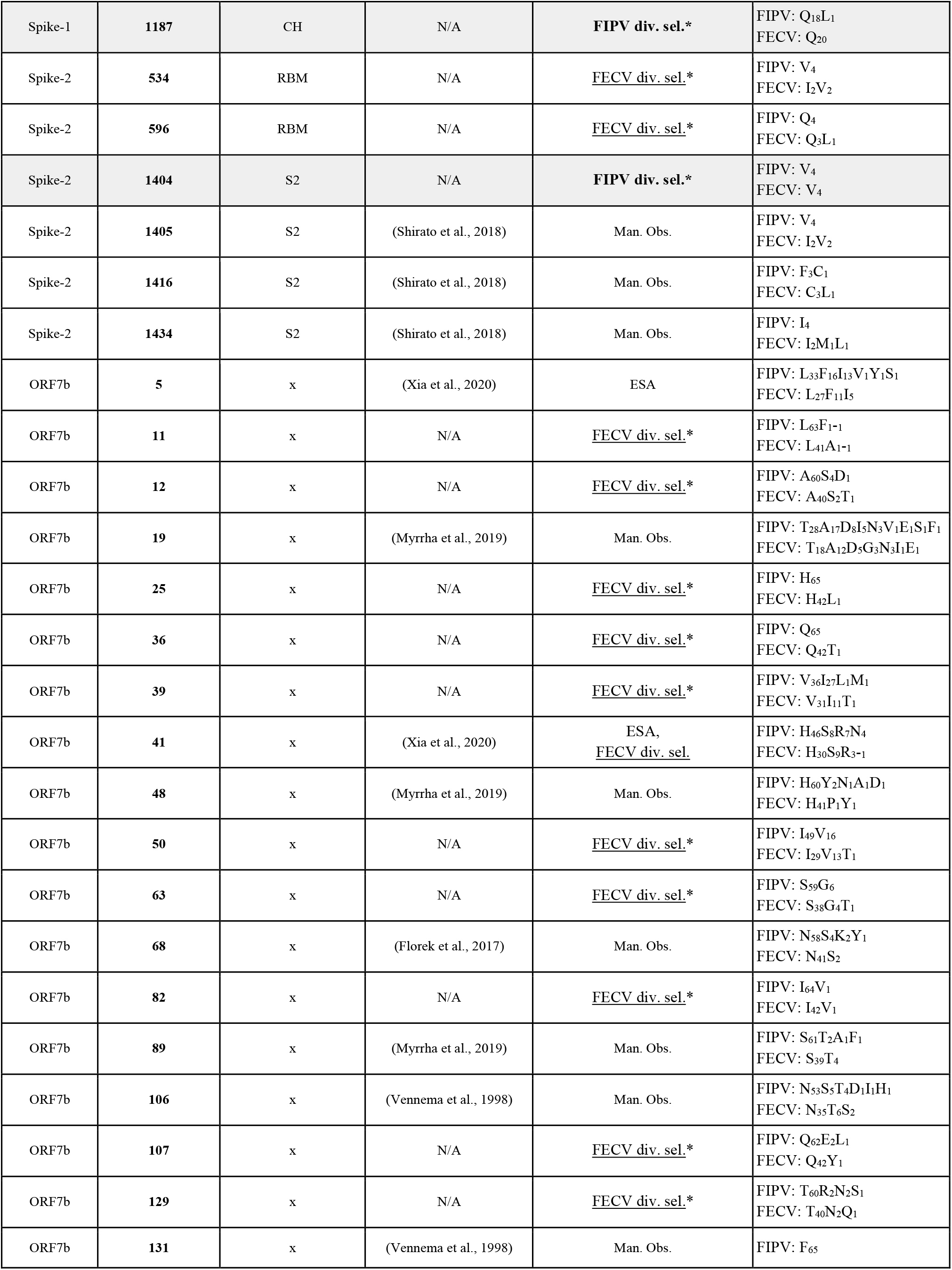
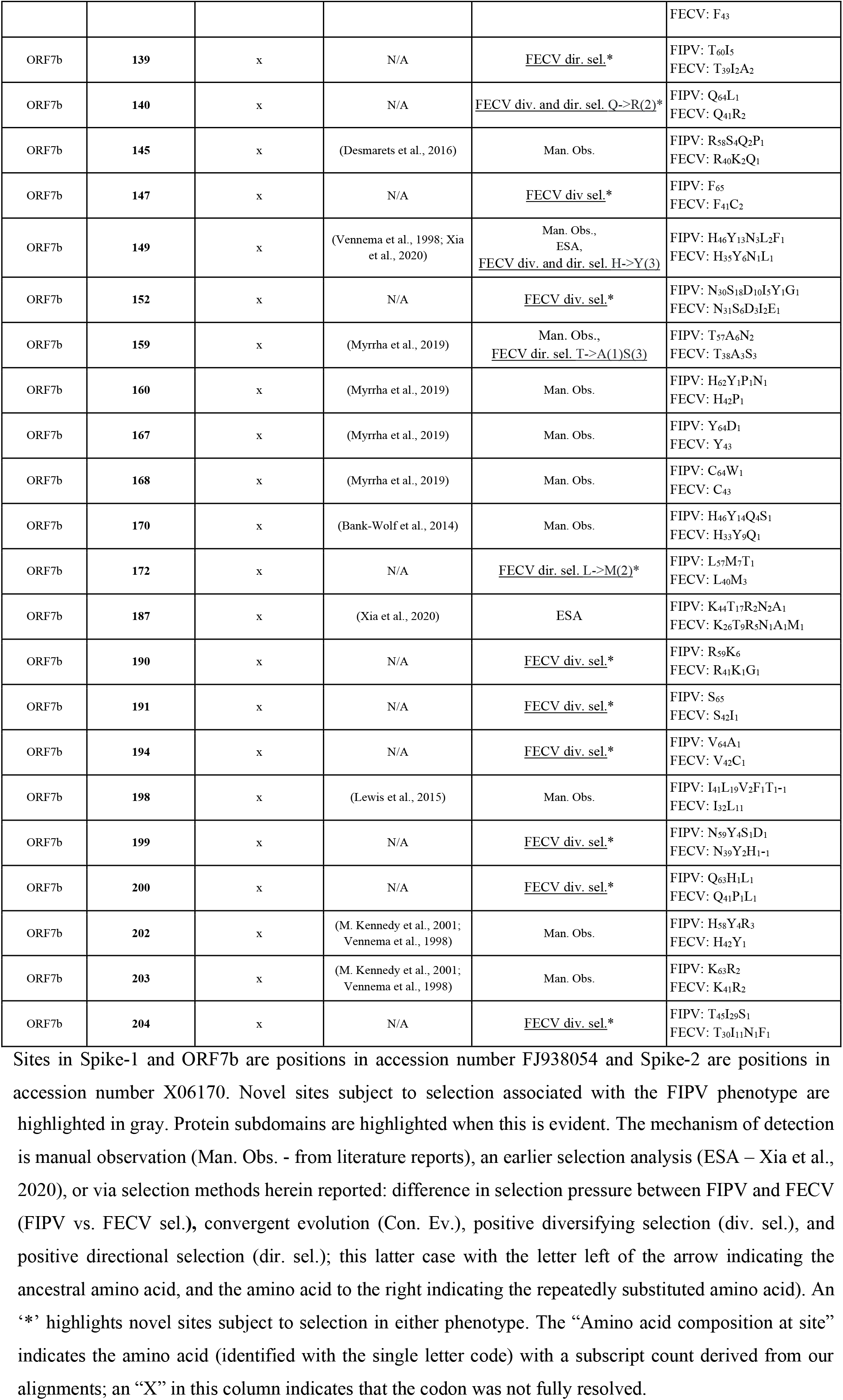
Sites identified to be subject to selection and/or manually observed, where selection is associated with either the FIPV, or FECV phenotype.

Relaxed selection in FIPV sequences relative to FECV sequences was identified in Spike-1 RFPs 7, 11, 12, and 13 (refer to Fig. 1 for functional subdomains included within each of those RFPs), and intensified selection in FIPV relative to FECV was identified in RFPs 8 and 9. A reduction in negative (purifying) selection in FIPV relative to FECV sequences was inferred in Spike-1 RFP 7, which encapsulates the S1/S2 FCS (**Fig. 2**). As a result, greater amino acid diversity can be observed in FIPV relative to FECV sequences for this region. Relaxed selection in FIPV sequences relative to FECV sequences was also identified in ORF3b and ORF7b (**Supplementary Table S4**).

**Figure 2.**
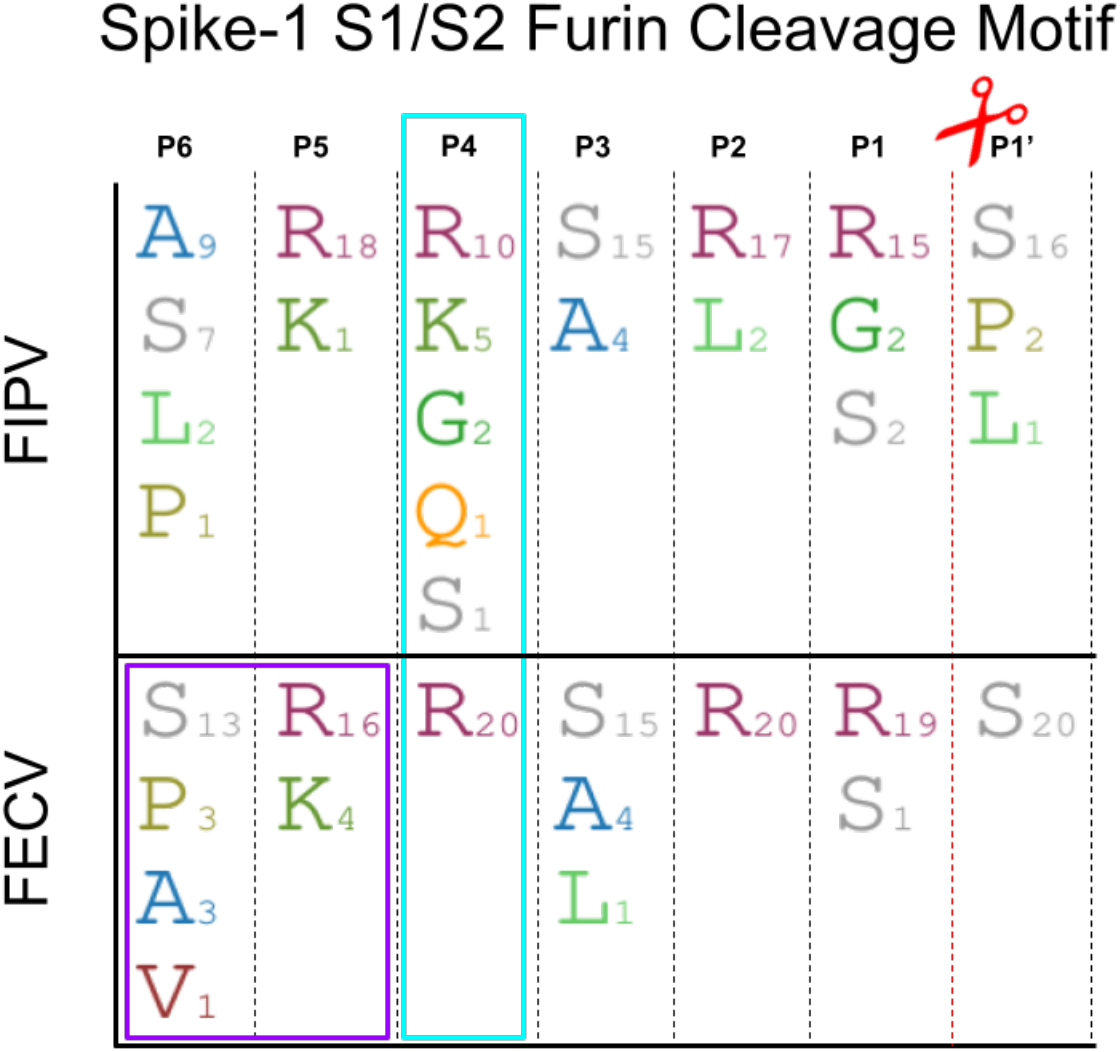
Spike-1 S1/S2 furin cleavage motif with the amino acid composition at critical sites involved in cleavage function (P6 to P1’ (Licitra et al., 2014)) for the FIPV and FECV sequences used in this study. The P6 and P5 sites were subject to directional selection in FECV sequences (highlighted in purple), and the P4 site was identified by the Contrast-FEL method (Kosakovsky Pond et al., 2021) (highlighted in cyan) to be evolving differently between the two phenotypes. Furin cleavage occurs between the P1 and P1’ site (Licitra et al., 2014), depicted with the red scissors.

The PCOC method (Rey et al., 2018) identified site 1058 within FIPV-1 Spike as evolving convergently (posterior probability > 0.8), and was the only site so identified. A Methionine (M) has been replaced with a Leucine (L) in the vast majority of FIPV sequences (**Fig. 3**).

**Figure 3.**
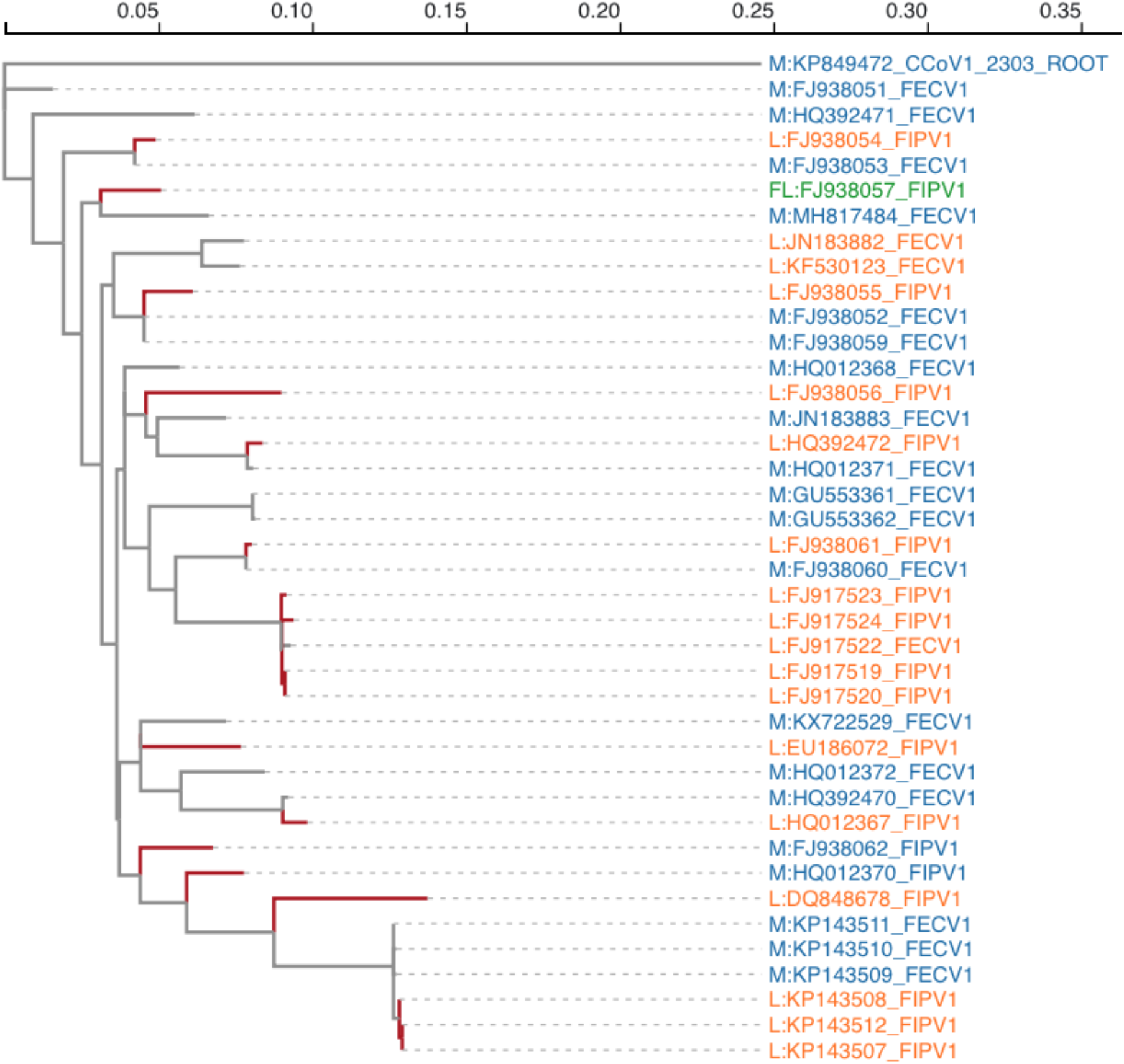
Convergent evolution detected at site 1058 within FIPV-1 Spike protein sequences by PCOC (Rey et al., 2018). Branches tested are highlighted in red. A Leucine (L) has arisen from a Methionine (M) in (15/18) FIPV sequences. Each leaf (tip) is annotated with the amino acid, accession number, and clinically diagnosed phenotype. The “FL” (FJ938057) represents an ambiguous base. CCoV-1 strain 23-03 Spike (KP849472) was used to root the tree.

### 3.4 Comparison of manually observed and selection inferred sites in FIPV and FECV

We compiled an extensive list of genetic mutations reported in the literature that differentiate FECV from FIPV sequences (**Table 2**). In instances where BUSTED-PH associated selection with the FIPV phenotype (Spike-1: RFPs 1, 5, 11, Spike-2: RFP 8) we identified 11 sites subject to selection not reported in the literature; 10 in Spike-1 and one in Spike-2 (**Table 2**), with the greatest concentration in the 0-domain of Spike-1 (**Fig. 1A**; **Table 2**). Out of the 46 sites either manually identified or reported in an earlier selection analysis (ESA) to differentiate FIPV from FECV sequences within RFPs associated with either FIPV or FECV positive selection, four were subject to positive selection associated with FIPV and two exhibited differences in selective regimes between FIPV and FECV sequences. The site most consistently reported in the literature differentiating FIPV from FECV *and* subject to positive selection was site 1058 (Bank-Wolf et al., 2014; Barker et al., 2017; H.-W. Chang et al., 2012; Decaro et al., 2021; Lewis et al., 2015; Ouyang et al., 2022; Xia et al., 2020) and our analyses suggest that this site also has a history of convergent evolution (**Fig. 3**). All sites inferred to be subject to positive selection in ORF7b were associated with the FECV phenotype, and of those 24 sites, only three have been previously identified in the literature (**Table 2**). Our analyses identified 38 sites under positive selection and associated with the FECV phenotype, of which 35 are previously unreported (**Table 2**). All other manually observed sites that fell within RFPs where BUSTED-PH could not distinguish selection signals associated with the FIPV or FECV phenotype are reported in **Supplementary Table S3**. Within ORF3c, a protein hypothesized to be involved in the shift in pathogenicity (Bank-Wolf et al., 2014; Borschensky & Reinacher, 2014; H.-W. Chang et al., 2010; Pedersen et al., 2012), only one of the six positively selected sites (site 165) has been previously identified in the literature (highlighted in **Supplementary Figure S1**). The FIPV phenotype was not uniquely associated with selection in ORF3c.

Sites in Spike-1 and ORF7b are positions in accession number FJ938054 and Spike-2 are positions in accession number X06170. Novel sites subject to selection associated with the FIPV phenotype are highlighted in gray. Protein subdomains are highlighted when this is evident. The mechanism of detection is manual observation (Man. Obs. - from literature reports), an earlier selection analysis (ESA – Xia et al., 2020), or via selection methods herein reported: difference in selection pressure between FIPV and FECV (FIPV vs. FECV sel.), convergent evolution (Con. Ev.), positive diversifying selection (div. sel.), and positive directional selection (dir. sel.); this latter case with the letter left of the arrow indicating the ancestral amino acid, and the amino acid to the right indicating the repeatedly substituted amino acid). An ‘*’ highlights novel sites subject to selection in either phenotype. The “Amino acid composition at site” indicates the amino acid (identified with the single letter code) with a subscript count derived from our alignments; an “X” in this column indicates that the codon was not fully resolved.

## 4. Discussion

Genetic mutations in FCoVs are linked to Feline Infectious Peritonitis (FIP) (Kipar & Meli, 2014) – an important infectious disease in wild felines and an often lethal disease in domestic felines worldwide. A shift in tropism, from epithelial cells (FECV) to macrophages/monocytes (FIPV), is associated with a subsequent shift in pathogenesis (Pedersen, 2014). While many genetic differences have been observed between FECV and FIPV sequences (**Table 2**), the specific phenotype-altering mutations within the FCoV genome remain unclear (M. A. Kennedy, 2020).

### 4.1 Cats as mixing vessels

Previous analyses examining natural selection differences between FECV and FIPV were methodologically limited, thus evolutionary genetic perspectives on putative phenotype-altering mutations remain largely unexplored. Felines are hub-species for a variety of coronavirus infections. Cats can be infected with both *Betacoronaviruses* (i.e., SARS-CoV and SARS-CoV-2 (Stout et al., 2020)) and *Alphacoronaviruses* (i.e, FCoVs, CCoV, TGEV, HCoV-229E (Li, 2016)), and there is convincing evidence to support a recombinant history between CCoV-2 and FCoV-1 (Herrewegh et al., 1998). Recent analyses of a newly discovered Canine Coronavirus isolated from symptomatic humans (CCoV-HuPn-2018; (Lednicky et al., 2022; Vlasova et al., 2022)), indicate a recombinant history involving FCoV-2 and CCoV-2 in the evolution of this new virus (Zehr et al., 2022) further highlighting the importance of felines as mixing vessels for CoVs. Therefore, it is of importance to identify where pathogenesis-altering mutations fall within FCoV genomes as an aid to interpreting how future recombination events might impact pathogenesis. Furthermore, information gained from these comparative evolutionary genetic analyses could be used to inform therapeutic strategies to combat infection, as well as gain a broader understanding of how evolutionary forces shape pathogenesis within FCoV genomes.

### 4.2 S1 Subunit

The S1 subunit of Spike in *Alphacoronaviruses* has been shown to play important functional roles in host cellular interactions (Li, 2015) and immune evasion (J. Shi et al., 2022; Y. Shi et al., 2021). Yang et al., (2020) reported on an extensive glycan repertoire across the S1 subunit of FIPV-1 Spike and suggested that virus entry, receptor recognition, and immune evasion may be impacted by this glycan shield. Similar glycan shielding functionality can be observed in the HIV-1 envelope protein, where heavy glycosylation on glycoprotein protein 120 (gp120) plays a crucial role in immune evasion (Pancera et al., 2014). Antibody-dependent enhancement (ADE), the process by which monoclonal antibodies (MAbs) enhance viral infection after binding, has been observed in FCoV-1 and -2 infections (Corapi et al., 1995; Hohdatsu et al., 1991; Olsen et al., 1992; Takano et al., 2008; Weiss & Scott, 1981), where the S1 subunit has been involved with ADE functionality (Takano et al., 2011). The majority of novel codon sites subject to positive selection and associated with the FIPV phenotype fall within the S1 subunit of the FCoV-1 Spike protein (five in 0 domain and two in B domain) (**Fig. 1A**). The S1 subunit comprises the NTD and CTD, where sugar binding and protein binding can occur, respectively (Li, 2016). The CTD of FIPV Spike-2 binds to fAPN during cellular entry, however, the principal FIPV-1 Spike receptor is not known (Dye et al., 2007; Hohdatsu et al., 1998; Tresnan et al., 1996). Recently, several Spike-1 receptors and attachment factors have been proposed, such as angiotensin converting enzyme-2 (ACE2) and dendritic cell-specific intercellular adhesion molecule grabbing non-integrin (DC-SIGN), respectively (Cook et al., 2022). Co-receptor and attachment factor binding for FIPV cannot be ruled out, as both lectins and carbohydrates, DC-SIGN and sialic acids, respectively, have been shown to interact with both Spike-1 and -2 (Cook et al., 2022; Desmarets et al., 2014; Regan et al., 2010; Regan & Whittaker, 2008). While CoV carbohydrate and proteinaceous receptor binding have been reported in the NTD and CTD respectively (Li, 2016), experimental studies will be necessary to confirm more precisely where in the FCoV Spike S1 subunit such binding may occur. Target cells for FIPV, macrophages/monocytes, contain sialoadhesin receptors on their cellular surface (O’Neill et al., 2013), which could interact with glycosylated sites on a fusion protein. Mutations within a specific region of the Spike 0-domain of a related *Alphacoronavirus-1*, TGEV, were shown to abrogate sialic acid co-receptor binding (Krempl et al., 1997, 2000; Schultze et al., 1996). Within the newly discovered CCoV-HuPn-2018 virus, Zehr et al., (2022) identified sites subject to positive selection in the homologous sialic acid binding region of CCoV-HuPn-2018, suggesting that adaptive change in sialic acid binding may have been relevant in the virus jump from dog to human. Here, in FIPV-1 Spike, we identify adaptively evolving sites in the 0-domain and suggest that this evolution may be associated with receptor binding functionality on target cells. Experimental studies will be necessary to verify if and where sialic acid binding occurs within the S1 subunit of FIPV-1 Spike, as well as to elucidate the functionality of sialic acid binding in FECV and FIPV infections (Cham et al., 2017; Desmarets et al., 2014).

Protein binding functionality is often contained within the CTD of the S1 subunit, which usually occurs within the RBD (Li, 2015). Y. Shi et al., (2021) examined the Spike structure of CoVs and remarked on the “lying” vs. “standing” (“up” vs. “down”, respectively) orientation of the RBD, where the *Alphacoronaviruses* studied had a “lying” or “down” RBD orientation. The group showed that an intact NTD from the Spike of HCoV-229E, an *Alphacoronavirus*, was essential for producing effective neutralizing antibodies (NAbs), compared to the Spike’s with a “standing” or “up” RBD that could generate effective NAbs from the RBD alone. Recently, a novel neutralizing epitope was identified in the NTD of HCoV-229E, where a single mutation in the NTD completely abolished NAb ability (J. Shi et al., 2022). The adaptation observed with the NTD of FIPV-1 could be associated with immune evasion, mirroring what has been shown in related *Alphacoronaviruses*. In the CTD, the two sites in FECV-2 Spike RBM (534 and 596) subject to positive selection fell within regions associated with adaptation to a new host in related *Alphacoronavirus-1s* (Olarte-Castillo et al., 2021). The sites subject to positive selection within the S1 subunit of Spike-1 and -2 may alter receptor recognition/ binding, facilitate immune evasion, and may even be associated with ADE; all processes that could contribute to FIP development.

### 4.3 Membrane Fusion

Membrane fusion takes place after receptor recognition and binding, and activation is a necessary step for class l viral fusion proteins to release the fusion peptide (FP). Activation can be accomplished in CoVs by a range of mechanisms – receptor binding, change in pH, and proteolytic cleavage (Bosch et al., 2003; Millet & Whittaker, 2015). There are two proteolytic cleavage motifs within the FCoV-1 Spike protein, the S1/S2 and S2’ sites, where the former is cleaved by furin (Licitra et al., 2013). Within the S1/S2 FCS, the composition at the P6, P4, P2, and P1 sites, specifically, having an arginine at each position, has been identified as critical for furin cleavage functionality, with an arginine residue at the P4 position being essential (Thomas, 2002). Of the two sites identified to be evolving under different selective regimes between FIPV and FECV sequences, one site, site 789, falls within the S1/S2 furin cleavage site (FCS) at the P4 position. We find that this position in FIPV sequences is under stronger diversifying, positive selection than in FECV sequences (see **Fig. 2**). Recent work from Ouyang et al., 2022 demonstrated that amino acid composition at the P4 site was highly diversified in FIPV sequences, while high amino acid conservation was observed in non-FIP sequences. Within *Betacoronaviruses*, such as Mouse Hepatitis Virus (MHV), a highly conserved P4 site (arginine) is also apparent (Stout et al., 2021). Within the S1/S2 FCS, relaxed selection (less purifying selection) was inferred in FIPV sequences relative to FECV sequences, further demonstrating the reduced evolutionary constraint at this location with FIPV sequences. Within FECV sequences, we identify directional selection at the P5 position from arginine towards lysine supporting the observation that an arginine at even sites within the S1/S2 FCS is favored. We did not identify detectable levels of positive selection uniquely associated with either phenotype at sites previously identified within the S2’ cleavage site of Spike to be associated with FIPV (Licitra et al., 2014). In related CoVs, Infectious Bronchitis Virus (IBV) and Human Coronavirus OC43 (HCoV-OC43), genetic mutations in the proteolytic cleavage sites in Spike were associated with alterations in tropism and pathogenesis (Belouzard et al., 2009; Le Coupanec et al., 2021; Tay et al., 2012; Yamada & Liu, 2009). The reduction in evolutionary constraint, coupled with abrogation of furin cleavage at the P4 site could suggest that furin cleavage functionality may not be critical to FIPV-1 Spike cellular entry. This is in contrast to FECV-1 Spike, where it appears that selection is shaping the FCS to be optimized. Perhaps, FIPV is using a furin cleavage-independent means of cellular entry, and may be using a co-receptor such as (fDC-SIGN or sialic acid). Importantly, it is encouraging that our hypothesis that the P4 site within the S1/S2 Spike-1 FCS may be putatively phenotype-altering is supported by newly collected data (Ouyang et al., 2022). Due to the conservative amino acid nature of this site in nonpathogenic sequences compared to the amino acid diversity observed in pathogenic sequences, this site may provide a useful diagnostic tool to identify FIPV sequences.

The Spike S2 subunit mediates membrane fusion and viral entry post-activation (Li, 2016). The fusion domain (FD) and heptad repeat regions 1 and 2 (HR1 and HR2, respectively) are hallmarks of class l virus fusion proteins that play a critical role in membrane fusion (Bosch et al., 2003). The FP within the FD inserts into the host cell membrane, and through the refolding process, a six helix bundle of HRs forms, ultimately resulting in the viral and cellular membranes being in close proximity (Bosch et al., 2003; White et al., 2008). The second site identified to be evolving measurably differently between FIPV and FECV sequences was canonical site 1058. First reported by H.-W. Chang et al., (2012), this site falls within the connecting region between the FD and HR1 and was the only site with detectable signals of convergent evolution in FIPV sequences (**Fig. 3**). More recently, Decaro et al., (2021) and Ouyang et al., (2022) reported similar findings to that of H.-W. Chang et al., (2012) with mutation M1058L observed in the vast majority of FIPV sequences. Mutation S1060A was also reported by H.-W. Chang et al., (2012) to differentiate FIPV from FECV sequences, but at this point does not appear to generalize to other data (Decaro et al., 2021; Ouyang et al., 2022) and is not subject to detectable signals of positive selection in our analysis. While mutation M1058L may not be a direct “switch” for phenotypic change (Barker et al., 2017; Jähne et al., 2022), the evidence of selection pressure acting on this site in FIPV sequences suggests that it may be involved in FIP development. Within FIPV-2 Spike we identified one novel site subject to positive selection in the S2 subunit in close proximity to positions identified by Rottier et al., (2005) in their mutation experiments involving amino acid positions within the HR1 and HR2 regions; these mutations inhibited macrophage entry (Rottier et al., 2005). Our results suggest that alterations in protein subdomains associated with membrane fusion may be associated with the development of FIP.

### 4.4 ORF3c and ORF7b

The association between genetic mutations in FCoV open reading frame 3c (ORF3c) and the FIPV phenotype has been the subject of considerable debate (Bank-Wolf et al., 2014; Borschensky & Reinacher, 2014; H.-W. Chang et al., 2010; Pedersen et al., 2012). Our analysis did not find selection within ORF3c to be associated with the FIPV phenotype. We did identify codon sites subject to positive selection within ORF3c of FCoV, of which, only one of the six positively selected sites has been previously identified in the literature (site 165) (**Supplementary Table S3**). *Betacoronaviruses* such as SARS-CoV-1 and -2 egress through lysosomal organelles, with ion channels of ORF3a from both viruses playing a critical role in this process (Ghosh et al., 2020; Kern et al., 2021; Lu et al., 2006). It has been shown that ORF3a from SARS-CoV-1 and FCoV ORF3c have similar predicted topologies (Oostra et al., 2006). Indeed, an alignment containing these two proteins suggests sequence homology (**Supplementary Figure S2**). The homologous site in SARS-CoV-2 ORF3a to FCoV ORF3c site 165 maps to a site critical for ion channel functionality (Kern et al., 2021) (**Supplementary Figure S1**). The hypothesis that FCoV ORF3c is a putative ion channel will need to be tested experimentally. Ion channels also play an important role in apoptosis (Lang et al., 2005), a phenomenon known to occur in FIPV infections (Haagmans et al., 1996; Shuid et al., 2015; Watanabe et al., 2018). It is possible that the adaptation we identify may be associated with viral egress and apoptosis from macrophages during an FCoV infection.

BUSTED-PH identified positive selection within ORF7b to be associated with the FECV phenotype, and a large number of sites (24) were identified from site-wise methods to be subject to positive selection. This ORF has been reported to be involved with ADE (Haijema et al., 2003), to interact with the Golgi retention signaling within the cell (Florek et al., 2017), and not be necessary for viral replication (Takano et al., 2011). Since FECV can be a chronic infection in the host (Herrewegh et al., 1997), and the host can be persistently infected with FECV (D. D. Addie et al., 2003; Kipar et al., 2010), the host immune system may act as a selective agent in FECV evolution. Our analysis identified relaxed selection within ORF7b in FIPV relative to FECV sequences, which could suggest an altered or diminished functional role of ORF7b in FIPV infections. In a related *Alphacoronavirus*, porcine respiratory coronavirus (PRCV), the loss of sialic acid binding functionality was associated with a large deletion in the NTD (Hulswit et al., 2016). We speculate that the adaptive evolution identified within this region may be associated with immune evasion in FECVs but that this functional role may not be necessary as FECV mutates to FIPV. Experimental studies will be necessary to interrogate sites under adaptive evolution in ORF7b to better understand their biological impact.

### 4.5 Diagnostic implications

Based on our results, there does not seem to be one or just a few mutations that define FIPV sequences, but rather, many, and the selection of so many sites within the host could be considered emblematic of short-sighted viral evolution. This, in turn, may contribute to the difficulty in identifying diagnostic sites in FCoV sequences, and the subsequent utility and reliability of such sites for an FIP diagnosis (Barker et al., 2017; Felten & Hartmann, 2019). Nonetheless, we report two sites subject to different selective regimes in FIPV and FECV sequences, as well as 11 novel sites subject to positive selection in FIPV sequences. A combination of sites reported herein may be needed to generate a “risk-score” assessment to aid in the diagnostic process (*e*.*g*., the more mutations identified, the higher the likelihood of FIP development). The majority of these sites fall within the NTD of FIPV-1 Spike, a protein subdomain associated with receptor recognition, receptor binding, and immune evasion in related *Alphacoronaviruses*; we hope this may provide a jumping-off point for future directed evolution experiments.

### 4.6 Limitations

There are limitations to this study. Specifically for Spike-2, selection signals identified may be limited by the relatively small number of sequences used, which can then impact the statistical confidence of parameter estimates and false positive rates. To account for this, we used methods that used a parametric bootstrap. Due to the reproducible and scalable nature of our computational methods and workflows, as more sequences become available data can be reanalyzed quickly. A field-wide, agreed-upon definition of an FECV sequence will also be useful in future comparative analyses.

## 5. Conclusion

In conclusion, we applied state-of-the-art comparative statistical methods to identify protein coding sites subject to positive selection pressure within FCoV genes previously hypothesized to be linked to the development of FIP. We found evidence of sites in Spike with an increased rate of positive selection in FIPV relative to FECV, as well as sites subject to positive selection associated with the FIPV phenotype that fell within protein subdomains associated with receptor binding and recognition, immune evasion, and membrane fusion. Perhaps, in the process of viral adaptation to evade host immune pressure and/or to escape the harsh gastrointestinal tract environment, the virus may acquire mutations that result in heightened virulence to the host, and ultimately, the increase in virulence could reduce the possibility of transmission – often referred to as short-sighted viral evolution (Lythgoe et al., 2017). We also report protein coding segments where relaxation of selection pressure is observed in FIPV relative to FECV that includes the S1/S2 FCS, which could suggest FIPV is using a furin-independent means of cellular entry. FIP is a complex disease, and it is likely that host factors contribute to disease onset beyond strictly viral factors (Borschensky & Reinacher, 2014), however, an animal model to propagate FCoV-1 virus *in vitro* remains to be established, making experimental validation difficult. Given the possible importance of host genetic variability and the development of FIP, we suggest a logical next step would be to examine FCoV quasispecies over the course of infection (Battilani et al., 2003; Desmarets et al., 2016; Gunn-Moore et al., 1999; Herrewegh et al., 1997; Hora et al., 2013; Kiss et al., 2000).

## Supporting information

SI-TableS3-SelectionResults

SI-TableS4-RELAXresults

SI-References

SI-FigureS1andS2captions

SI-TableS1-AccessionsUsed

SI-TableS2-RecombinantBreakpoints

SI-FigureS1-3DstructureORF3c

SI-FigureS2-ProteinAlignment

## 6. Acknowledgements

We would like to thank everyone in the Temple Viral Evolution Group, along with <u>Alyssa Pivirotto</u>, Amanda Wilson, and Avery Selberg who all offered valuable editorial suggestions throughout the writing process.

## 7. Data Availability

All data used herein are publicly available on GenBank. Accession numbers used can be found in Supplementary Table S1.

