## Supplementary material for "Natural selection differences detected in key protein domains between non-pathogenic and pathogenic Feline Coronavirus phenotypes": SI-TableS3-SelectionResults

**Supplementary Table S3**. Sites identified to be subject to selection and/or manually observed, where selection is not solely associated with either the FIPV, or FECV phenotype.

| Protein | Position in reference | Protein subdomain | Relevant literature | Identification method | Amino acid composition at site |
|---|---|---|---|---|---|
| Spike-1 | **117** | 0 | (Lewis et al., 2015) | Man. Obs. | FIPV: $T_8A_5P_3Q_2V_1$<br>FECV: $A_6T_6L_2E_1Q_1D_1I_1P_1X_1$ |
| Spike-1 | **118** | 0 | (Desmarets et al., 2016) | Man. Obs. | FIPV: $R_{13}I_1Q_1A_1V_1S_1C_1$<br>FECV: $R_{12}Y_2H_2M_1A_1V_1G_1$ |
| Spike-1 | **130** | 0 | N/A | FCoV dir. sel. N->H(1)T(1), T->K(2)N(1)* | FIPV: $N_8K_6T_4Q_1$<br>FECV: $K_9T_7N_1H_1S_1Q_1$ |
| Spike-1 | **134** | 0 | N/A | FCoV dir. sel. D->S(1)* | FIPV: $S_{19}$<br>FECV: $S_{20}$ |
| Spike-1 | **142** | 0 | (Lewis et al., 2015) | Man. Obs. | FIPV: $G_8D_4N_3E_2K_1A_1$<br>FECV: $D_7E_4N_4G_3R_1S_1$ |
| Spike-1 | **144** | 0 | N/A | FCoV div. sel.* | FIPV: $E_{18}Q_1$<br>FECV: $E_{16}Y_2Q_1V_1$ |
| Spike-1 | **145** | 0 | N/A | FCoV dir. sel. E->A(5)* | FIPV: $E_{13}A_5D_1$<br>FECV: $E_{11}A_9$ |
| Spike-1 | **148** | 0 | N/A | FCoV dir. sel. L->P(1)* | FIPV: $P_{19}$<br>FECV: $P_{20}$ |
| Spike-1 | **149** | 0 | N/A | FCoV dir. sel. H->D(2)* | FIPV: $H_{18}D_1$<br>FECV: $H_{16}D_3Y_1$ |
| Spike-1 | **158** | 0 | (Desmarets et al., 2016) | Man. Obs. | FIPV: $K_{11}Q_7N_1$<br>FECV: $K_{14}Q_2R_2N_1P_1$ |
| Spike-1 | **162** | 0 | N/A | FCoV dir. sel. S->I(1)* | FIPV: $I_{19}$<br>FECV: $I_{20}$ |
| Spike-1 | **165** | 0 | N/A | FCoV div. sel.* | FIPV: $Y_{16}I_3$<br>FECV: $Y_{15}I_3H_2$ |
| Spike-1 | **166** | 0 | N/A | FCoV dir. sel. N->H(1)* | FIPV: $H_{19}$<br>FECV: $H_{20}$ |
| Spike-1 | **167** | 0 | (Desmarets et al., 2016) | Man. Obs. | FIPV: $Q_{10}A_5K_2E_1R_1$<br>FECV: $Q_{15}E_4R_1$ |
| Spike-1 | **169** | 0 | N/A | FCoV div. sel.* | FIPV: $S_7T_5N_2D_1G_1A_1K_1R_1$<br>FECV: $S_{11}N_3D_2G_1K_1R_1T_1$ |
| Spike-1 | **177** | 0 | N/A | FCoV dir. sel. D->Q(1)* | FIPV: $Q_{18}Y_1$<br>FECV: $Q_{19}L_1$ |
| Spike-1 | **219** | 0 | N/A | FCoV div. sel.* | FIPV: $E_{10}D_5N_2V_1G_1$<br>FECV: $D_9E_8N_2V_1$ |
| Spike-1 | **221** | 0 | N/A | FCoV dir. sel. T->D(1)* | FIPV: $D_{18}K_1$<br>FECV: $D_{20}$ |
| Spike-1 | **226** | 0 | N/A | FCoV dir. sel. D->E(1)* | FIPV: $E_{19}$<br>FECV: $E_{20}$ |
| Spike-1 | **227** | 0 | N/A | FCoV dir. sel. A->S(1), S->A(2), V->A(1)* | FIPV: $A_{13}S_6$<br>FECV: $A_{11}S_9$ |
| Spike-1 | **229** | 0 | N/A | FCoV dir. sel.* | FIPV: $I_{18}M_1$<br>FECV: $I_{20}$ |
| Spike-1 | **230** | 0 | N/A | FCoV dir. sel.* | FIPV: $S_{19}$<br>FECV: $S_{20}$ |

| | | | | | |
|---|---|---|---|---|---|
| Spike-1 | **234** | 0 | (Lewis et al., 2015) | Man. Obs. | FIPV: $N_{14}S_5$<br>FECV: $N_{12}S_8$ |
| Spike-1 | **235** | 0 | (Desmarets et al., 2016; Lewis et al., 2015) | Man. Obs. | FIPV: $Q_6K_6R_4A_1G_1N_1$<br>FECV: $R_5Q_4S_3H_2X_2A_1T_1G_1K_1$ |
| Spike-1 | **236** | 0 | N/A | FCoV div. and dir. sel. I->A(1)L(2)Y(3)* | FIPV: $I_{11}L_5T_1Y_1N_1$<br>FECV: $I_9Y_5L_4A_1V_1$ |
| Spike-1 | **238** | 0 | N/A | FCoV div. sel.* | FIPV: $Y_{18}S_1$<br>FECV: $Y_{14}Q_2K_1S_1R_1X_1$ |
| Spike-1 | **255** | 0 | N/A | FCoV dir. sel. N->S(3)* | FIPV: $N_{10}S_9$<br>FECV: $N_{14}S_6$ |
| Spike-1 | **271** | 0 | N/A | FCoV div. sel.* | FIPV: $S_{16}T_2D_1$<br>FECV: $S_{19}T_1$ |
| Spike-1 | **292** | A | N/A | FCoV dir. sel. S->G(2)* | FIPV: $S_{14}G_4X_1$<br>FECV: $S_{15}G_5$ |
| Spike-1 | **293** | A | (Licitra et al., 2013) | Man. Obs. | FIPV: $F_{18}Y_1$<br>FECV: $F_{20}$ |
| Spike-1 | **335** | A | (Desmarets et al., 2016) | Man. Obs. | FIPV: $V_{19}$<br>FECV: $V_{20}$ |
| Spike-1 | **339** | A | N/A | FCoV div. sel.* | FIPV: $S_{11}K_4N_4$<br>FECV: $S_6N_5K_4D_3T_2$ |
| Spike-1 | **350** | A | N/A | FCoV dir. sel. D->E(3)* | FIPV: $D_{10}E_9$<br>FECV: $D_{13}E_7$ |
| Spike-1 | **362** | A | N/A | FCoV dir. sel. A->D(1)* | FIPV: $D_{18}A_1$<br>FECV: $D_{20}$ |
| Spike-1 | **371** | A | (Vennema et al., 1998) | Man. Obs. | FIPV: $D_{17}E_1G_1$<br>FECV: $D_{12}E_7G_1$ |
| Spike-1 | **372** | A | (Lewis et al., 2015) | Man. Obs., FCoV div. sel. | FIPV: $Q_{16}R_3$<br>FECV: $Q_{18}R_2$ |
| Spike-1 | **384** | A | N/A | FCoV div. sel.* | FIPV: $S_{16}G_3$<br>FECV: $S_{15}G_3T_1A_1$ |
| Spike-1 | **396** | A | N/A | FCoV div. sel.* | FIPV: $F_{10}N_3Y_3\text{-}_2L_1$<br>FECV: $F_{12}N_4Y_3Q_1$ |
| Spike-1 | **399** | A | N/A | FCoV div. sel.* | FIPV: $A_{13}N_3V_1T_1\text{-}_1$<br>FECV: $A_{14}N_4T_1V_1$ |
| Spike-1 | **401** | A | N/A | FCoV div. sel.* | FIPV: $N_{14}Q_2G_1S_1\text{-}_1$<br>FECV: $N_{17}Q_2G_1$ |
| Spike-1 | **404** | A | N/A | FCoV div. sel.* | FIPV: $S_{16}A_2\text{-}_1$<br>FECV: $S_{20}$ |
| Spike-1 | **406** | A | (Lewis et al., 2015) | Man. Obs. | FIPV: $I_{10}V_5T_2S_1\text{-}_1$<br>FECV: $I_{20}$ |
| Spike-1 | **410** | A | N/A | FCoV div. sel.* | FIPV: $K_{15}S_2R_1\text{-}_1$<br>FECV: $K_{18}R_1S_1$ |
| Spike-1 | **GAP (410-411)** | A | N/A | FCoV div. sel.* | FIPV: $M_6T_4N_2D_2\text{-}_2S_1R_1A_1$<br>FECV: $M_6T_6N_2E_2D_2Q_1A_1$ |
| Spike-1 | **411** | A | N/A | FCoV div. sel.* | FIPV: $H_{13}N_2Y_2Q_1\text{-}_1$<br>FECV: $H_{13}N_2Y_2S_1Q_1C_1$ |
| Spike-1 | **422** | A | N/A | FCoV div. sel.* | FIPV: $S_{12}N_5A_1\text{-}_1$<br>FECV: $S_{19}A_1$ |

| | | | | | |
|---|---|---|---|---|---|
| Spike-1 | **423** | A | N/A | FCoV div. sel.* | FIPV: $I_6V_6H_3S_1P_1L_1\text{-}_1$<br>FECV: $V_{11}I_4H_3T_1M_1$ |
| Spike-1 | **435** | A | (Lewis et al., 2015) | Man. Obs.,<br>FCoV div. sel. | FIPV: $T_{18}I_1$<br>FECV: $T_{19}I_1$ |
| Spike-1 | **456** | A | N/A | FCoV div. sel.* | FIPV: $S_{18}M_1$<br>FECV: $S_{18}M_1N_1$ |
| Spike-1 | **458** | A | N/A | FCoV div. sel.* | FIPV: $P_{15}Q_4$<br>FECV: $P_{15}Q_3S_2$ |
| Spike-1 | **472** | A | N/A | FCoV div. and dir. sel.<br>N->K(1)Q(2), Q->N(1)* | FIPV: $N_{15}Q_3K_1$<br>FECV: $N_{13}Q_5K_1H_1$ |
| Spike-1 | **493** | A | (Vennema et al., 1998) | Man. Obs. | FIPV: $T_{18}A_1$<br>FECV: $T_{20}$ |
| Spike-1 | **505** | A | N/A | FCoV dir. sel. L->I(2)* | FIPV: $L_{12}I_7$<br>FECV: $I_{10}L_{10}$ |
| Spike-1 | **647** | B | N/A | FCoV div. sel.* | FIPV: $V_{16}T_3$<br>FECV: $V_{15}T_4L_1$ |
| Spike-1 | **648** | B | N/A | FCoV dir. sel. N->S(2)* | FIPV: $S_{10}N_9$<br>FECV: $N_{15}S_5$ |
| Spike-1 | **663** | B | (Desmarets et al., 2016;<br>Lewis et al., 2015) | Man. Obs. | FIPV: $K_{13}R_3N_2T_1$<br>FECV: $K_{11}R_4M_3I_1T_1$ |
| Spike-1 | **844** | S1/S2 | (Xia et al., 2020) | ESA,<br>FCoV div. sel.* | FIPV: $H_{16}K_3$<br>FECV: $K_{10}H_9R_1$ |
| Spike-1 | **846** | S1/S2 | N/A | FCoV dir. sel. E->A(2)* | FIPV: $E_{14}A_5$<br>FECV: $E_{18}A_2$ |
| Spike-1 | **847** | S1/S2 | N/A | FCoV dir. sel. T->I(1)* | FIPV: $I_{15}T_4$<br>FECV: $I_{17}T_2N_1$ |
| Spike-1 | **851** | S1/S2 | (Vennema et al., 1998) | Man. Obs. | FIPV: $S_{12}G_5N_1\text{-}_1$<br>FECV: $S_{17}T_1N_1G_1$ |
| Spike-1 | **864** | S1/S2 | (Lewis et al., 2015) | Man. Obs. | FIPV: $S_{13}T_3F_2L_1$<br>FECV: $S_{13}T_7$ |
| Spike-1 | **902** | UH | (Xia et al., 2020) | ESA | FIPV: $N_8T_6S_5$<br>FECV: $N_9T_7S_2K_2$ |
| Spike-1 | **905** | UH | N/A | FCoV dir. sel. T->A(2)* | FIPV: $T_{13}A_5K_1$<br>FECV: $T_{14}A_4S_2$ |
| Spike-1 | **919** | UH | (Desmarets et al., 2016) | Man. Obs. | FIPV: $N_{18}S_1$<br>FECV: $N_{20}$ |
| Spike-1 | **950** | S2’ | (Xia et al., 2020) | ESA | FIPV: $T_{14}N_2A_2S_1$<br>FECV: $T_{13}S_3N_2A_1K_1$ |
| Spike-1 | **952** | S2’ | (Xia et al., 2020) | ESA | FIPV: $V_{13}A_4L_1D_1$<br>FECV: $V_9A_5T_3L_1I_1X_1$ |
| Spike-1 | **953** | S2' | N/A | FCoV dir. sel. V->L(2)* | FIPV: $L_{10}V_8T_1$<br>FECV: $L_{13}V_7$ |
| Spike-1 | **954** | S2' | N/A | FCoV div. sel.* | FIPV: $G_{17}R_2$<br>FECV: $G_{20}$ |
| Spike-1 | **967** | S2' | (Xia et al., 2020) | ESA | FIPV: $R_{11}S_6K_2$<br>FECV: $R_{11}S_9$ |
| Spike-1 | **968** | S2' | (Xia et al., 2020) | ESA,<br>FCoV div. sel.* | FIPV: $S_7Y_3E_3A_2D_2V_1H_1$<br>FECV: $S_9E_5D_3N_3$ |

| | | | | | |
|---|---|---|---|---|---|
| Spike-1 | **973** | S2' | (Xia et al., 2020) | ESA,<br>FCoV div. sel.* | FIPV: $R_9T_6K_2S_2$<br>FECV: $T_9R_4S_3K_2G_1V_1$ |
| Spike-1 | **976** | S2’ | (Xia et al., 2020) | ESA | FIPV: $K_{14}M_3V_2$<br>FECV: $K_{16}R_3T_1$ |
| Spike-1 | **1030** | FP | (Vennema et al., 1998;<br>Xia et al., 2020) | Man. Obs.,<br>ESA,<br>FCoV div. sel. | FIPV: $Q_6D_4G_3H_3Y_1A_1X_1$<br>FECV: $D_{11}Q_7G_1S_1$ |
| Spike-1 | **1276** | CD | (Xia et al., 2020) | ESA | FIPV: $N_{12}G_7$<br>FECV: $G_{13}N_7$ |
| Spike-1 | **1313** | CD | N/A | FCoV dir. sel. T->K(2)* | FIPV: $T_{12}K_5N_1M_1$<br>FECV: $T_{14}K_3S_1A_1I_1$ |
| Spike-1 | **1342** | HR2 | (Lewis et al., 2015) | Man. Obs. | FIPV: $P_{12}S_5L_1X_1$<br>FECV: $P_{18}S_2$ |
| Spike-1 | **1378** | HR2 | (Lewis et al., 2015) | Man. Obs. | FIPV: $E_{16}Q_2S_1$<br>FECV: $E_{20}$ |
| Spike-2 | **223** | S1 | N/A | FCoV div. sel.* | FIPV: $Y_3K_1$<br>FECV: $H_3K_1$ |
| Spike-2 | **938** | S2 | (Shirato et al., 2018) | Man. Obs. | FIPV: $S_2N_1T_1$<br>FECV: $N_4$ |
| Spike-2 | **961** | S2 | (Shirato et al., 2018) | Man. Obs. | FIPV: $G_3R_1$<br>FECV: $R_4$ |
| Spike-2 | **972** | S2 | (Shirato et al., 2018) | Man. Obs. | FIPV: $V_4$<br>FECV: $V_4$ |
| Spike-2 | **1014** | S2 | (Rottier et al., 2005;<br>Shirato et al., 2018) | Man. Obs. | FIPV: $A_3D_1$<br>FECV: $D_4$ |
| Spike-2 | **1094** | S2 | (Shirato et al., 2018) | Man. Obs. | FIPV: $Q_4$<br>FECV: $Q_3K_1$ |
| Spike-2 | **1237** | S2 | (Shirato et al., 2018) | Man. Obs. | FIPV: $S_4$<br>FECV: $S_3P_1$ |
| Spike-2 | **1279** | S2 | (Shirato et al., 2018) | Man. Obs. | FIPV: $V_3A_1$<br>FECV: $A_3V_1$ |
| Spike-2 | **1333** | S2 | (Shirato et al., 2018) | Man. Obs. | FIPV: $F_3L_1$<br>FECV: $L_3F_1$ |
| ORF3a | **30** | x | N/A | FCoV dir. sel. E->L(1)* | FIPV: $L_{42}V_1$<br>FECV: $L_{38}$ |
| ORF3a | **32** | x | N/A | FCoV dir. sel. K->T(1),<br>T->I(1)N(2)* | FIPV: $N_{22}T_{20}I_1$<br>FECV: $N_{19}T_{16}I_2K_1$ |
| ORF3a | **47** | x | N/A | FCoV div. sel.* | FIPV: $E_{41}Q_1D_1$<br>FECV: $E_{31}H_7$ |
| ORF3a | **58** | x | N/A | FCoV dir. sel. G->Q(1)* | FIPV: $Q_{40}H_2L_1$<br>FECV: $Q_{38}$ |
| ORF3a | **61-62 GAP** | x | N/A | FCoV dir. sel. V->E(1)* | FIPV: $E_{41}A_1{-}_1$<br>FECV: $E_{38}$ |
| ORF3a | **64** | x | N/A | FCoV div. sel.* | FIPV: $P_{27}S_8F_6L_1R_1$<br>FECV: $P_{22}S_8F_3L_3H_2$ |
| ORF3a | **65** | x | N/A | FCoV div. sel.* | FIPV: $N_{40}D_2T_1$<br>FECV: $N_{34}D_4$ |
| ORF3b | **2** | x | N/A | FCoV div. sel.* | FIPV: $P_{30}L_2$<br>FECV: $P_{26}L_1$ |

| | | | | | |
|---|---|---|---|---|---|
| ORF3b | **64** | x | N/A | FCoV div. sel.* | FIPV: $K_{19}R_{13}$<br>FECV: $K_{14}R_{13}$ |
| ORF3b | **71** | x | N/A | FCoV div. sel.* | FIPV: $A_{24}S_4L_2T_1E_1$<br>FECV: $A_{19}S_4V_2T_1L_1$ |
| ORF3c | **11** | x | N/A | FCoV dir. sel. S->G(3)* | FIPV: $S_{24}G_3$<br>FECV: $S_{37}G_{11}R_1$ |
| ORF3c | **71** | x | N/A | FCoV div. sel.* | FIPV: $G_{24}S_3$<br>FECV: $G_{34}S_{15}$ |
| ORF3c | **72** | x | N/A | FCoV div. sel.* | FIPV: $V_{26}I_1$<br>FECV: $V_{48}I_1$ |
| ORF3c | **159** | x | N/A | FCoV div. sel.* | FIPV: $M_{27}$<br>FECV: $M_{46}T_2I_1$ |
| ORF3c | **165** | 190 in SC2 3a | (Lewis et al., 2015) | Man. Obs.,<br>FCoV div. sel. | FIPV: $T_{23}M_4$<br>FECV: $T_{49}$ |
| ORF3c | **175** | x | N/A | FCoV div. and dir. sel.<br>G->C(2)* | FIPV: $G_{27}$<br>FECV: $G_{44}C_5$ |
| ORF7a | **6** | x | (Lewis et al., 2015) | Man. Obs. | FIPV: $H_{41}Y_1$<br>FECV: $H_{21}Y_1$ |
| ORF7a | **9** | x | (Bank-Wolf et al., 2014) | Man. Obs. | FIPV: $L_{29}F_{12}I_1$<br>FECV: $L_{12}F_{10}$ |
| ORF7a | **12** | x | (Bank-Wolf et al., 2014) | Man. Obs. | FIPV: $V_{23}A_{19}$<br>FECV: $V_{13}A_9$ |
| ORF7a | **47** | x | (Bank-Wolf et al., 2014) | Man. Obs. | FIPV: $S_{26}N_{15}I_1$<br>FECV: $S_{19}N_2T_1$ |
| ORF7a | **69** | x | N/A | FCoV div. sel.* | FIPV: $K_{39}R_3$<br>FECV: $K_{20}R_2$ |

Spike-1, ORF3c, ORF7a mapped to FJ938054 -- FIPV strain UU4 "Black", Spike-2 mapped to X06170 – FIPV strain 79-1146, ORF3a mapped to KJ665842 -- FIPV, ORF3b mapped to EU664276 -- FIPV. Protein subdomains are highlighted when this is evident. The mechanism of detection is manual observation (Man. Obs. - from literature reports), an earlier selection analysis (ESA – Xia et al., 2020), or via selection methods herein reported: difference in selection pressure between FIPV and FECV (FIPV vs. FECV sel.), positive diversifying selection (div. sel.), and positive directional selection (dir. sel.); this latter case with the letter left of the arrow indicating the ancestral amino acid, and the amino acid to the right indicating the repeatedly substituted amino acid). An '*' highlights novel sites subject to selection herein identified. The "Amino acid composition at site" indicates the amino acid (identified with the single letter code) with a subscript count derived from our alignments; an "X" in this column indicates that the codon was not fully resolved.
