## Supplementary material for "Natural selection differences detected in key protein domains between non-pathogenic and pathogenic Feline Coronavirus phenotypes": SI-TableS4-RELAXresults

**Supplementary Table S4**. RELAX results indicating either intensification or relaxation of selection in FIPV relative to FECV

| Protein | Recombinant Free Partition (RFP) | Relaxation or Intesification | p-value | Likelihood ratio (LR) |
|---|---|---|---|---|
| Spike-1 | 7 | Relaxation (K = 0.56) | 0 | 19.18 |
| Spike-1 | 8 | Intensification (K = 14.84) | 0.019 | 5.46 |
| Spike-1 | 9 | Intensification (K = 6.24) | 0.003 | 8.6 |
| Spike-1 | 11 | Relaxation (K = 0.68) | 0.002 | 9.53 |
| Spike-1 | 12 | Relaxation (K = 0.0) | 0.01 | 6.69 |
| Spike-1 | 13 | Relaxation (K = 0.58) | 0.013 | 6.16 |
| ORF3b | 1 | Relaxation (K = 0.17) | 0.016 | 5.85 |
| ORF7b | 1 | Relaxation (K = 0.64) | 0 | 12.71 |
