## Supplementary material for "Natural selection differences detected in key protein domains between non-pathogenic and pathogenic Feline Coronavirus phenotypes": SI-References

### Supplementary Material References


Bank-Wolf, B. R., Stallkamp, I., Wiese, S., Moritz, A., Tekes, G., & Thiel, H.-J. (2014). Mutations of 3c and spike protein genes correlate with the occurrence of feline infectious peritonitis. *Veterinary Microbiology*, *173*(3–4), 177–188. https://doi.org/10.1016/j.vetmic.2014.07.020

Desmarets, L. M. B., Vermeulen, B. L., Theuns, S., Conceição-Neto, N., Zeller, M., Roukaerts, I. D. M., Acar, D. D., Olyslaegers, D. A. J., Van Ranst, M., Matthijnssens, J., & Nauwynck, H. J. (2016). Experimental feline enteric coronavirus infection reveals an aberrant infection pattern and shedding of mutants with impaired infectivity in enterocyte cultures. *Scientific Reports*, *6*, 20022. https://doi.org/10.1038/srep20022

Kern, D. M., Sorum, B., Mali, S. S., Hoel, C. M., Sridharan, S., Remis, J. P., Toso, D. B., Kotecha, A., Bautista, D. M., & Brohawn, S. G. (2021). Cryo-EM structure of SARS-CoV-2 ORF3a in lipid nanodiscs. *Nature Structural & Molecular Biology*, *28*(7), 573–582. https://doi.org/10.1038/s41594-021-00619-0

Lewis, C. S., Porter, E., Matthews, D., Kipar, A., Tasker, S., Helps, C. R., & Siddell, S. G. (2015). Genotyping coronaviruses associated with feline infectious peritonitis. *The Journal of General Virology*, *96*(Pt 6), 1358–1368. https://doi.org/10.1099/vir.0.000084

Licitra, B. N., Millet, J. K., Regan, A. D., Hamilton, B. S., Rinaldi, V. D., Duhamel, G. E., & Whittaker, G. R. (2013). Mutation in spike protein cleavage site and pathogenesis of feline coronavirus. *Emerging Infectious Diseases*, *19*(7), 1066–1073. https://doi.org/10.3201/eid1907.121094

Rottier, P. J. M., Nakamura, K., Schellen, P., Volders, H., & Haijema, B. J. (2005). Acquisition of macrophage tropism during the pathogenesis of feline infectious peritonitis is determined by mutations in the feline coronavirus spike protein. *Journal of Virology*, *79*(22), 14122–14130. https://doi.org/10.1128/JVI.79.22.14122-14130.2005

Shirato, K., Chang, H.-W., & Rottier, P. J. M. (2018). Differential susceptibility of macrophages to serotype II feline coronaviruses correlates with differences in the viral spike protein. *Virus Research*, *255*, 14–23. https://doi.org/10.1016/j.virusres.2018.06.010

Vennema, H., Poland, A., Foley, J., & Pedersen, N. C. (1998). Feline infectious peritonitis viruses arise by mutation from endemic feline enteric coronaviruses. *Virology*, *243*(1), 150–157. https://doi.org/10.1006/viro.1998.9045

Xia, H., Li, X., Zhao, W., Jia, S., Zhang, X., Irwin, D. M., & Zhang, S. (2020). Adaptive Evolution of Feline Coronavirus Genes Based on Selection Analysis. *BioMed Research International*, *2020*, 9089768. https://doi.org/10.1155/2020/9089768
