## Supplementary material for "Natural selection differences detected in key protein domains between non-pathogenic and pathogenic Feline Coronavirus phenotypes": SI-FigureS1andS2captions

**Supplementary Figure S1. Structural modeling of FCoV-1 UU4 ORF3c using AlphaFold 2**. The amino acid sequence encoding the ORF3c protein of FCoV-1 UU4 (accession no. FJ938054.1) was used to model its structure as a homodimer with AlphaFold 2 Multimer and ColabFold software (https://colab.research.google.com/github/sokrypton/ColabFold/blob/main/AlphaFold2.ipynb). N-terminal and C-terminal regions with low confidence scores (pLDDT) are not displayed. For comparison, the cryo-EM-determined structure of SARS-CoV-2 ORF3a is shown (right panel). The residue at position 165 of FCoV-1 UU4 ORF3c is highlighted in red (T165), as well as the corresponding residue in SARS-CoV-2 ORF3a (T190). The amino acid composition at this site in FIPV sequences was 23 Threonine's and 4 Methionine's; this site was conserved for Threonine's in FECV sequences.

**Supplementary Figure S2. Protein alignment of ORF3a Beta- and ORF3c Alphacoronaviruses**. SARS-CoV-2 ORF3a is used as a reference (highlighted as "REF") and FIPV strain UU4 is highlighted with "FIPV". Secondary structures are highlighted at indices where appropriate. The lower tunnel of SARS-CoV-2 ORF3a putative ion channel identified by Kern et al., 2021 is highlighted in green.
