## Supplementary material for "Natural selection differences detected in key protein domains between non-pathogenic and pathogenic Feline Coronavirus phenotypes": SI-TableS1-AccessionsUsed

**Supplementary Table S1**. All Accessions used in this study

| Protein coding sequence | Accession number | Biotype and Serotype |
|---|---|---|
| ORF3a | EU664276 | FIPV1 |
| ORF3a | EU664277 | FECV1 |
| ORF3a | EU664278 | FIPV1 |
| ORF3a | EU664279 | FECV1 |
| ORF3a | EU664280 | FECV1 |
| ORF3a | EU664283 | FECV1 |
| ORF3a | EU664288 | FECV1 |
| ORF3a | EU664291 | FIPV1 |
| ORF3a | EU664293 | FECV1 |
| ORF3a | EU664296 | FIPV1 |
| ORF3a | EU664297 | FIPV1 |
| ORF3a | FJ917519 | FIPV1 |
| ORF3a | FJ917520 | FIPV1 |
| ORF3a | FJ917521 | FIPV1 |
| ORF3a | FJ917522 | FECV1 |
| ORF3a | FJ917523 | FIPV1 |
| ORF3a | FJ917524 | FIPV1 |
| ORF3a | FJ917525 | FIPV1 |
| ORF3a | FJ917526 | FIPV1 |
| ORF3a | FJ917527 | FIPV1 |
| ORF3a | FJ917528 | FIPV1 |
| ORF3a | FJ917529 | FIPV1 |
| ORF3a | FJ938051 | FECV1 |
| ORF3a | FJ938052 | FECV1 |
| ORF3a | FJ938053 | FECV1 |
| ORF3a | FJ938055 | FIPV1 |
| ORF3a | FJ938056 | FIPV1 |
| ORF3a | FJ938057 | FIPV1 |
| ORF3a | FJ938058 | FIPV1 |
| ORF3a | FJ938059 | FECV1 |

| | | |
|---|---|---|
| ORF3a | FJ938060 | FECV1 |
| ORF3a | FJ938061 | FIPV1 |
| ORF3a | FJ938062 | FIPV1 |
| ORF3a | FJ943761 | FECV1 |
| ORF3a | FJ943762 | FECV1 |
| ORF3a | FJ943764 | FECV1 |
| ORF3a | FJ943771 | FECV1 |
| ORF3a | GU553361 | FECV1 |
| ORF3a | GU553362 | FECV1 |
| ORF3a | HQ012367 | FIPV1 |
| ORF3a | HQ012368 | FECV1 |
| ORF3a | HQ012369 | FIPV1 |
| ORF3a | HQ012370 | FIPV1 |
| ORF3a | HQ012371 | FECV1 |
| ORF3a | HQ012372 | FECV1 |
| ORF3a | HQ392470 | FECV1 |
| ORF3a | HQ392471 | FECV1 |
| ORF3a | HQ392472 | FIPV1 |
| ORF3a | JN183882 | FECV1 |
| ORF3a | JN183883 | FECV1 |
| ORF3a | KF530123 | FECV1 |
| ORF3a | KJ665813 | FECV1 |
| ORF3a | KJ665814 | FECV1 |
| ORF3a | KJ665815 | FECV1 |
| ORF3a | KJ665816 | FECV1 |
| ORF3a | KJ665817 | FECV1 |
| ORF3a | KJ665818 | FECV1 |
| ORF3a | KJ665819 | FECV1 |
| ORF3a | KJ665820 | FECV1 |
| ORF3a | KJ665821 | FECV1 |
| ORF3a | KJ665822 | FECV1 |

| | | |
|---|---|---|
| ORF3a | KJ665823 | FECV1 |
| ORF3a | KJ665824 | FECV1 |
| ORF3a | KJ665825 | FECV1 |
| ORF3a | KJ665826 | FECV1 |
| ORF3a | KJ665827 | FECV1 |
| ORF3a | KJ665828 | FECV1 |
| ORF3a | KJ665829 | FECV1 |
| ORF3a | KJ665830 | FECV1 |
| ORF3a | KJ665831 | FECV1 |
| ORF3a | KJ665832 | FECV1 |
| ORF3a | KJ665833 | FIPV1 |
| ORF3a | KJ665834 | FIPV1 |
| ORF3a | KJ665835 | FIPV1 |
| ORF3a | KJ665836 | FIPV1 |
| ORF3a | KJ665837 | FIPV1 |
| ORF3a | KJ665838 | FIPV1 |
| ORF3a | KJ665839 | FIPV1 |
| ORF3a | KJ665840 | FIPV1 |
| ORF3a | KJ665841 | FIPV1 |
| ORF3a | KJ665842 | FIPV1 |
| ORF3a | KJ665843 | FIPV1 |
| ORF3a | KJ665844 | FIPV1 |
| ORF3a | KJ665845 | FIPV1 |
| ORF3a | KJ665846 | FIPV1 |
| ORF3a | KJ665850 | FIPV1 |
| ORF3a | KJ665851 | FIPV1 |
| ORF3a | KJ665852 | FIPV1 |
| ORF3a | KJ665853 | FIPV1 |
| ORF3a | KJ665854 | FIPV1 |
| ORF3a | KJ665855 | FIPV1 |
| ORF3a | KJ665856 | FIPV1 |

| | | |
|---|---|---|
| ORF3a | KJ665857 | FIPV1 |
| ORF3a | KJ665858 | FIPV1 |
| ORF3a | KJ665859 | FIPV1 |
| ORF3a | KJ665860 | FIPV1 |
| ORF3a | KJ665861 | FIPV1 |
| ORF3a | KP143507 | FIPV1 |
| ORF3a | KP143508 | FIPV1 |
| ORF3a | KP143509 | FECV1 |
| ORF3a | KP143510 | FECV1 |
| ORF3a | KP143511 | FECV1 |
| ORF3a | KP143512 | FIPV1 |
| ORF3a | KX722529 | FECV1 |
| ORF3a | MH817484 | FECV1 |
| ORF3a | MT444152 | FIPV1 |
| ORF3a | MW545831 | FECV1 |
| ORF3a | MW545832 | FECV1 |
| ORF3a | MW545833 | FECV1 |
| ORF3a | MW545835 | FECV1 |
| ORF3a | MW545836 | FECV1 |
| ORF3a | MW545837 | FECV1 |
| ORF3a | MW545838 | FECV1 |
| ORF3a | MW545839 | FIPV1 |
| ORF3a | MW545840 | FECV1 |
| ORF3a | MW545841 | FIPV1 |
| ORF3a | MW545842 | FECV1 |
| ORF3b | EU664276 | FIPV1 |
| ORF3b | EU664277 | FECV1 |
| ORF3b | EU664278 | FIPV1 |
| ORF3b | EU664279 | FECV1 |
| ORF3b | EU664280 | FECV1 |
| ORF3b | EU664283 | FECV1 |

| | | |
|---|---|---|
| ORF3b | EU664288 | FECV1 |
| ORF3b | EU664291 | FIPV1 |
| ORF3b | EU664296 | FIPV1 |
| ORF3b | FJ917519 | FIPV1 |
| ORF3b | FJ917520 | FIPV1 |
| ORF3b | FJ917521 | FIPV1 |
| ORF3b | FJ917522 | FECV1 |
| ORF3b | FJ917523 | FIPV1 |
| ORF3b | FJ917524 | FIPV1 |
| ORF3b | FJ917526 | FIPV1 |
| ORF3b | FJ917528 | FIPV1 |
| ORF3b | FJ917529 | FIPV1 |
| ORF3b | FJ938051 | FECV1 |
| ORF3b | FJ938052 | FECV1 |
| ORF3b | FJ938053 | FECV1 |
| ORF3b | FJ938055 | FIPV1 |
| ORF3b | FJ938056 | FIPV1 |
| ORF3b | FJ938057 | FIPV1 |
| ORF3b | FJ938058 | FIPV1 |
| ORF3b | FJ938060 | FECV1 |
| ORF3b | FJ938059 | FECV1 |
| ORF3b | FJ938062 | FIPV1 |
| ORF3b | FJ943761 | FECV1 |
| ORF3b | FJ943762 | FECV1 |
| ORF3b | FJ943764 | FECV1 |
| ORF3b | FJ943771 | FECV1 |
| ORF3b | GU553361 | FECV1 |
| ORF3b | GU553362 | FECV1 |
| ORF3b | HQ012368 | FECV1 |
| ORF3b | HQ012369 | FIPV1 |
| ORF3b | HQ012370 | FIPV1 |

| | | |
|---|---|---|
| ORF3b | HQ012371 | FECV1 |
| ORF3b | HQ012372 | FECV1 |
| ORF3b | HQ392470 | FECV1 |
| ORF3b | HQ392471 | FECV1 |
| ORF3b | HQ392472 | FIPV1 |
| ORF3b | JN183882 | FECV1 |
| ORF3b | JN183883 | FECV1 |
| ORF3b | KF530123 | FECV1 |
| ORF3b | KJ665814 | FECV1 |
| ORF3b | KJ665817 | FECV1 |
| ORF3b | KJ665820 | FECV1 |
| ORF3b | KJ665824 | FECV1 |
| ORF3b | KJ665825 | FECV1 |
| ORF3b | KJ665827 | FECV1 |
| ORF3b | KJ665828 | FECV1 |
| ORF3b | KJ665829 | FECV1 |
| ORF3b | KJ665830 | FECV1 |
| ORF3b | KJ665835 | FIPV1 |
| ORF3b | KJ665836 | FIPV1 |
| ORF3b | KJ665837 | FIPV1 |
| ORF3b | KJ665838 | FIPV1 |
| ORF3b | KJ665839 | FIPV1 |
| ORF3b | KJ665840 | FIPV1 |
| ORF3b | KJ665843 | FIPV1 |
| ORF3b | KJ665844 | FIPV1 |
| ORF3b | KJ665845 | FIPV1 |
| ORF3b | KJ665847 | FIPV1 |
| ORF3b | KJ665848 | FIPV1 |
| ORF3b | KJ665852 | FIPV1 |
| ORF3b | KJ665853 | FIPV1 |
| ORF3b | KJ665854 | FIPV1 |

| | | |
|---|---|---|
| ORF3b | KJ665855 | FIPV1 |
| ORF3b | KJ665856 | FIPV1 |
| ORF3b | KJ665860 | FIPV1 |
| ORF3b | KP143507 | FIPV1 |
| ORF3b | KP143508 | FIPV1 |
| ORF3b | KP143509 | FECV1 |
| ORF3b | KP143510 | FECV1 |
| ORF3b | KP143511 | FECV1 |
| ORF3b | KP143512 | FIPV1 |
| ORF3b | KX722529 | FECV1 |
| ORF3b | MH817484 | FECV1 |
| ORF3b | MW545868 | FECV1 |
| ORF3b | MW545870 | FIPV1 |
| ORF3b | MW545871 | FECV1 |
| ORF3b | MW545872 | FIPV1 |
| ORF3b | MW545873 | FECV1 |
| ORF3c | FJ917522 | FECV1 |
| ORF3c | FJ938051 | FECV1 |
| ORF3c | FJ938052 | FECV1 |
| ORF3c | FJ938053 | FECV1 |
| ORF3c | FJ938054 | FIPV1 |
| ORF3c | FJ938055 | FIPV1 |
| ORF3c | FJ938059 | FECV1 |
| ORF3c | FJ938060 | FECV1 |
| ORF3c | FJ943761 | FECV1 |
| ORF3c | FJ943762 | FECV1 |
| ORF3c | FJ943763 | FECV1 |
| ORF3c | FJ943771 | FECV1 |
| ORF3c | GU053607 | FIPV1 |
| ORF3c | GU053608 | FIPV1 |
| ORF3c | GU053609 | FIPV1 |

| | | |
|---|---|---|
| ORF3c | GU053611 | FIPV1 |
| ORF3c | GU053612 | FIPV1 |
| ORF3c | GU053613 | FECV1 |
| ORF3c | GU053614 | FECV1 |
| ORF3c | GU053615 | FECV1 |
| ORF3c | GU053616 | FECV1 |
| ORF3c | GU053618 | FECV1 |
| ORF3c | GU053619 | FECV1 |
| ORF3c | GU053621 | FECV1 |
| ORF3c | GU053622 | FECV1 |
| ORF3c | GU053623 | FECV1 |
| ORF3c | GU053624 | FECV1 |
| ORF3c | GU053625 | FECV1 |
| ORF3c | GU053626 | FECV1 |
| ORF3c | GU053627 | FECV1 |
| ORF3c | GU053628 | FECV1 |
| ORF3c | GU053629 | FECV1 |
| ORF3c | GU053630 | FECV1 |
| ORF3c | GU053632 | FECV1 |
| ORF3c | GU053635 | FECV1 |
| ORF3c | GU053636 | FECV1 |
| ORF3c | GU053637 | FECV1 |
| ORF3c | GU053638 | FECV1 |
| ORF3c | GU053639 | FECV1 |
| ORF3c | GU053644 | FIPV1 |
| ORF3c | GU053646 | FIPV1 |
| ORF3c | GU053648 | FIPV1 |
| ORF3c | GU053651 | FIPV1 |
| ORF3c | GU053653 | FIPV1 |
| ORF3c | GU053656 | FIPV1 |
| ORF3c | GU053660 | FIPV1 |

| | | |
|---|---|---|
| ORF3c | GU053663 | FIPV1 |
| ORF3c | GU053666 | FIPV1 |
| ORF3c | GU553361 | FECV1 |
| ORF3c | GU553362 | FECV1 |
| ORF3c | HQ012368 | FECV1 |
| ORF3c | HQ012371 | FECV1 |
| ORF3c | HQ012372 | FECV1 |
| ORF3c | HQ392470 | FECV1 |
| ORF3c | HQ392471 | FECV1 |
| ORF3c | JN183883 | FECV1 |
| ORF3c | KF530123 | FECV1 |
| ORF3c | KJ665813 | FECV1 |
| ORF3c | KJ665815 | FECV1 |
| ORF3c | KJ665817 | FECV1 |
| ORF3c | KJ665818 | FECV1 |
| ORF3c | KJ665819 | FECV1 |
| ORF3c | KJ665820 | FECV1 |
| ORF3c | KJ665821 | FECV1 |
| ORF3c | KJ665822 | FECV1 |
| ORF3c | KJ665823 | FECV1 |
| ORF3c | KJ665824 | FECV1 |
| ORF3c | KJ665825 | FECV1 |
| ORF3c | KJ665826 | FECV1 |
| ORF3c | KJ665827 | FECV1 |
| ORF3c | KJ665828 | FECV1 |
| ORF3c | KJ665829 | FECV1 |
| ORF3c | KJ665830 | FECV1 |
| ORF3c | KJ665832 | FECV1 |
| ORF3c | KJ665833 | FIPV1 |
| ORF3c | KJ665837 | FIPV1 |
| ORF3c | KJ665838 | FIPV1 |

| | | |
|---|---|---|
| ORF3c | KJ665840 | FIPV1 |
| ORF3c | KJ665847 | FIPV1 |
| ORF3c | KJ665848 | FIPV1 |
| ORF3c | KJ665849 | FIPV1 |
| ORF3c | KJ665852 | FIPV1 |
| ORF3c | KJ665853 | FIPV1 |
| ORF3c | KJ665855 | FIPV1 |
| ORF3c | KJ665857 | FIPV1 |
| ORF3c | KJ665861 | FIPV1 |
| ORF3c | KP143508 | FIPV1 |
| ORF3c | KP143509 | FECV1 |
| ORF3c | KP143510 | FECV1 |
| ORF3c | KP143511 | FECV1 |
| ORF3c | KX722529 | FECV1 |
| ORF3c | MH817484 | FECV1 |
| ORF7a | AY994055 | FIPV2 |
| ORF7a | DQ010921 | FIPV2 |
| ORF7a | DQ848678 | FIPV1 |
| ORF7a | EU186072 | FIPV1 |
| ORF7a | FJ917519 | FIPV1 |
| ORF7a | FJ917520 | FIPV1 |
| ORF7a | FJ917521 | FIPV1 |
| ORF7a | FJ917522 | FECV1 |
| ORF7a | FJ917523 | FIPV1 |
| ORF7a | FJ917524 | FIPV1 |
| ORF7a | FJ917525 | FIPV1 |
| ORF7a | FJ917526 | FIPV1 |
| ORF7a | FJ917527 | FIPV1 |
| ORF7a | FJ917528 | FIPV1 |
| ORF7a | FJ917529 | FIPV1 |
| ORF7a | FJ938051 | FECV1 |

| | | |
|---|---|---|
| ORF7a | FJ938052 | FECV1 |
| ORF7a | FJ938053 | FECV1 |
| ORF7a | FJ938054 | FIPV1 |
| ORF7a | FJ938055 | FIPV1 |
| ORF7a | FJ938056 | FIPV1 |
| ORF7a | FJ938057 | FIPV1 |
| ORF7a | FJ938058 | FIPV1 |
| ORF7a | FJ938059 | FECV1 |
| ORF7a | FJ938060 | FECV1 |
| ORF7a | FJ938061 | FIPV1 |
| ORF7a | FJ938062 | FIPV1 |
| ORF7a | FJ943761 | FECV1 |
| ORF7a | FJ943763 | FECV1 |
| ORF7a | FJ943764 | FECV1 |
| ORF7a | GU553361 | FECV1 |
| ORF7a | GU553362 | FECV1 |
| ORF7a | HQ012368 | FECV1 |
| ORF7a | HQ012369 | FIPV1 |
| ORF7a | HQ012370 | FIPV1 |
| ORF7a | HQ012371 | FECV1 |
| ORF7a | HQ012372 | FECV1 |
| ORF7a | HQ392470 | FECV1 |
| ORF7a | HQ392471 | FECV1 |
| ORF7a | HQ392472 | FIPV1 |
| ORF7a | JN183882 | FECV1 |
| ORF7a | JN183883 | FECV1 |
| ORF7a | JN634064 | FIPV1 |
| ORF7a | KF530123 | FECV1 |
| ORF7a | KJ665779 | FECV1 |
| ORF7a | KJ665780 | FECV1 |
| ORF7a | KJ665781 | FECV1 |

| | | |
|---|---|---|
| ORF7a | KJ665782 | FECV1 |
| ORF7a | KJ665783 | FECV1 |
| ORF7a | KJ665784 | FECV1 |
| ORF7a | KJ665785 | FECV1 |
| ORF7a | KJ665786 | FECV1 |
| ORF7a | KJ665787 | FIPV1 |
| ORF7a | KJ665788 | FIPV1 |
| ORF7a | KJ665789 | FIPV1 |
| ORF7a | KJ665790 | FIPV1 |
| ORF7a | KJ665791 | FIPV1 |
| ORF7a | KJ665792 | FIPV1 |
| ORF7a | KJ665793 | FIPV1 |
| ORF7a | KJ665794 | FIPV1 |
| ORF7a | KJ665795 | FIPV1 |
| ORF7a | KJ665796 | FIPV1 |
| ORF7a | KJ665797 | FIPV1 |
| ORF7a | KJ665798 | FIPV1 |
| ORF7a | KJ665799 | FIPV1 |
| ORF7a | KJ665800 | FIPV1 |
| ORF7a | KJ665801 | FIPV1 |
| ORF7a | KP143507 | FIPV1 |
| ORF7a | KP143508 | FIPV1 |
| ORF7a | KP143509 | FECV1 |
| ORF7a | KP143510 | FECV1 |
| ORF7a | KP143511 | FECV1 |
| ORF7a | KP143512 | FIPV1 |
| ORF7a | KX722529 | FECV1 |
| ORF7a | MH817484 | FECV1 |
| ORF7a | MT444152 | FIPV1 |
| ORF7a | NC_002306 | FIPV2 |
| ORF7a | X90570 | FIPV |

| | | |
|---|---|---|
| ORF7a | X90571 | FIPV |
| ORF7a | X90572 | FIPV |
| ORF7a | X90573 | FIPV |
| ORF7a | X90574 | FECV |
| ORF7a | X90575 | FIPV |
| ORF7a | X90576 | FIPV |
| ORF7a | X90577 | FIPV |
| ORF7a | X90578 | FIPV |
| ORF7b | AY994055 | FIPV2 |
| ORF7b | DQ010921 | FIPV2 |
| ORF7b | DQ648122 | FIP |
| ORF7b | DQ675414 | FIP |
| ORF7b | DQ675417 | FIP |
| ORF7b | DQ675418 | FIP |
| ORF7b | DQ675419 | FIP |
| ORF7b | DQ675420 | FECV |
| ORF7b | DQ675421 | FECV |
| ORF7b | DQ675422 | FECV |
| ORF7b | DQ675423 | FECV |
| ORF7b | DQ675428 | FIP |
| ORF7b | DQ675429 | FIP |
| ORF7b | DQ675430 | FIP |
| ORF7b | DQ675431 | FIP |
| ORF7b | DQ675432 | FIP |
| ORF7b | DQ675433 | FECV |
| ORF7b | DQ675434 | FECV |
| ORF7b | DQ675436 | FIP |
| ORF7b | DQ675437 | FECV |
| ORF7b | DQ675438 | FECV |
| ORF7b | DQ675439 | FECV |
| ORF7b | DQ675440 | FIP |

| | | |
|---|---|---|
| ORF7b | DQ675441 | FECV |
| ORF7b | DQ675442 | FECV |
| ORF7b | DQ675443 | FECV |
| ORF7b | DQ675444 | FECV |
| ORF7b | DQ675445 | FIP |
| ORF7b | DQ675446 | FIP |
| ORF7b | DQ675447 | FECV |
| ORF7b | DQ675448 | FIP |
| ORF7b | DQ675449 | FIP |
| ORF7b | DQ675450 | FIP |
| ORF7b | DQ675451 | FECV |
| ORF7b | DQ675452 | FIP |
| ORF7b | DQ848678 | FIPV1 |
| ORF7b | FJ917519 | FIPV1 |
| ORF7b | FJ917520 | FIPV1 |
| ORF7b | FJ917521 | FIPV1 |
| ORF7b | FJ917522 | FECV1 |
| ORF7b | FJ917523 | FIPV1 |
| ORF7b | FJ917524 | FIPV1 |
| ORF7b | FJ917525 | FIPV1 |
| ORF7b | FJ917526 | FIPV1 |
| ORF7b | FJ917527 | FIPV1 |
| ORF7b | FJ917528 | FIPV1 |
| ORF7b | FJ917529 | FIPV1 |
| ORF7b | FJ938051 | FECV1 |
| ORF7b | FJ938052 | FECV1 |
| ORF7b | FJ938053 | FECV1 |
| ORF7b | FJ938054 | FIPV1 |
| ORF7b | FJ938055 | FIPV1 |
| ORF7b | FJ938056 | FIPV1 |
| ORF7b | FJ938057 | FIPV1 |

| | | |
|---|---|---|
| ORF7b | FJ938058 | FIPV1 |
| ORF7b | FJ938059 | FECV1 |
| ORF7b | FJ938060 | FECV1 |
| ORF7b | FJ938061 | FIPV1 |
| ORF7b | FJ938062 | FIPV1 |
| ORF7b | FJ943761 | FECV1 |
| ORF7b | FJ943763 | FECV1 |
| ORF7b | FJ943764 | FECV1 |
| ORF7b | GU553361 | FECV1 |
| ORF7b | GU553362 | FECV1 |
| ORF7b | HQ012367 | FIPV1 |
| ORF7b | HQ012368 | FECV1 |
| ORF7b | HQ012369 | FIPV1 |
| ORF7b | HQ012370 | FIPV1 |
| ORF7b | HQ012371 | FECV1 |
| ORF7b | HQ012372 | FECV1 |
| ORF7b | HQ392470 | FECV1 |
| ORF7b | HQ392471 | FECV1 |
| ORF7b | HQ392472 | FIPV1 |
| ORF7b | JN183882 | FECV1 |
| ORF7b | JN183883 | FECV1 |
| ORF7b | JX239089 | FECV |
| ORF7b | JX239090 | FECV |
| ORF7b | JX239091 | FIPV |
| ORF7b | JX239092 | FIPV |
| ORF7b | JX239093 | FECV |
| ORF7b | JX239094 | FECV |
| ORF7b | JX239095 | FIPV |
| ORF7b | JX239096 | FECV |
| ORF7b | JX239097 | FECV |
| ORF7b | JX239098 | FIPV |

| | | |
|---|---|---|
| ORF7b | JX239099 | FECV |
| ORF7b | JX239100 | FECV |
| ORF7b | JX239101 | FECV |
| ORF7b | JX239102 | FECV |
| ORF7b | JX239103 | FIPV |
| ORF7b | JX239104 | FIPV |
| ORF7b | JX239105 | FIPV |
| ORF7b | JX239106 | FIPV |
| ORF7b | JX239107 | FECV |
| ORF7b | JX239108 | FECV |
| ORF7b | JX239109 | FIPV |
| ORF7b | JX239110 | FECV |
| ORF7b | JX239111 | FIPV |
| ORF7b | JX239112 | FIPV |
| ORF7b | JX239113 | FECV |
| ORF7b | JX239114 | FECV |
| ORF7b | JX239115 | FIPV |
| ORF7b | JX239116 | FECV |
| ORF7b | JX239117 | FECV |
| ORF7b | JX239118 | FECV |
| ORF7b | JX239119 | FECV |
| ORF7b | JX239120 | FIPV |
| ORF7b | JX239121 | FECV |
| ORF7b | JX239122 | FECV |
| ORF7b | JX239123 | FIPV |
| ORF7b | JX239124 | FECV |
| ORF7b | JX239125 | FECV |
| ORF7b | JX239126 | FIPV |
| ORF7b | JX239127 | FECV |
| ORF7b | JX239128 | FECV |
| ORF7b | JX239129 | FECV |

| | | |
|---|---|---|
| ORF7b | JX239130 | FECV |
| ORF7b | JX239131 | FECV |
| ORF7b | JX239132 | FECV |
| ORF7b | JX239133 | FIPV |
| ORF7b | JX239134 | FECV |
| ORF7b | JX239135 | FECV |
| ORF7b | JX239136 | FECV |
| ORF7b | JX239137 | FECV |
| ORF7b | JX239138 | FECV |
| ORF7b | JX239139 | FECV |
| ORF7b | KF530123 | FECV1 |
| ORF7b | KJ665779 | FECV1 |
| ORF7b | KJ665780 | FECV1 |
| ORF7b | KJ665781 | FECV1 |
| ORF7b | KJ665782 | FECV1 |
| ORF7b | KJ665783 | FECV1 |
| ORF7b | KJ665784 | FECV1 |
| ORF7b | KJ665785 | FECV1 |
| ORF7b | KJ665786 | FECV1 |
| ORF7b | KJ665787 | FIPV1 |
| ORF7b | KJ665788 | FIPV1 |
| ORF7b | KJ665789 | FIPV1 |
| ORF7b | KJ665790 | FIPV1 |
| ORF7b | KJ665791 | FIPV1 |
| ORF7b | KJ665793 | FIPV1 |
| ORF7b | KJ665794 | FIPV1 |
| ORF7b | KJ665795 | FIPV1 |
| ORF7b | KJ665796 | FIPV1 |
| ORF7b | KJ665797 | FIPV1 |
| ORF7b | KJ665798 | FIPV1 |
| ORF7b | KJ665799 | FIPV1 |

| | | |
|---|---|---|
| ORF7b | KJ665800 | FIPV1 |
| ORF7b | KJ665801 | FIPV1 |
| ORF7b | KP143507 | FIPV1 |
| ORF7b | KP143508 | FIPV1 |
| ORF7b | KP143509 | FECV1 |
| ORF7b | KP143510 | FECV1 |
| ORF7b | KP143511 | FECV1 |
| ORF7b | KP143512 | FIPV1 |
| ORF7b | KX722529 | FECV1 |
| ORF7b | MH817484 | FECV1 |
| ORF7b | NC_002306 | FIPV2 |
| ORF7b | X90570 | FIPV |
| ORF7b | X90571 | FIPV |
| ORF7b | X90572 | FIPV |
| ORF7b | X90573 | FIPV |
| ORF7b | X90574 | FECV |
| ORF7b | X90575 | FIPV |
| ORF7b | X90577 | FIPV |
| ORF7b | X90578 | FIPV |
| Spike-1 | DQ848678 | FIPV1 |
| Spike-1 | EU186072 | FIPV1 |
| Spike-1 | FJ917519 | FIPV1 |
| Spike-1 | FJ917520 | FIPV1 |
| Spike-1 | FJ917521 | FIPV1 |
| Spike-1 | FJ917522 | FECV1 |
| Spike-1 | FJ917523 | FIPV1 |
| Spike-1 | FJ917524 | FIPV1 |
| Spike-1 | FJ938051 | FECV1 |
| Spike-1 | FJ938052 | FECV1 |
| Spike-1 | FJ938053 | FECV1 |
| Spike-1 | FJ938054 | FIPV1 |

| | | |
|---|---|---|
| Spike-1 | FJ938055 | FIPV1 |
| Spike-1 | FJ938056 | FIPV1 |
| Spike-1 | FJ938057 | FIPV1 |
| Spike-1 | FJ938059 | FECV1 |
| Spike-1 | FJ938060 | FECV1 |
| Spike-1 | FJ938061 | FIPV1 |
| Spike-1 | FJ938062 | FIPV1 |
| Spike-1 | GU553361 | FECV1 |
| Spike-1 | GU553362 | FECV1 |
| Spike-1 | HQ012367 | FIPV1 |
| Spike-1 | HQ012368 | FECV1 |
| Spike-1 | HQ012370 | FIPV1 |
| Spike-1 | HQ012371 | FECV1 |
| Spike-1 | HQ012372 | FECV1 |
| Spike-1 | HQ392470 | FECV1 |
| Spike-1 | HQ392471 | FECV1 |
| Spike-1 | HQ392472 | FIPV1 |
| Spike-1 | JN183882 | FECV1 |
| Spike-1 | JN183883 | FECV1 |
| Spike-1 | KF530123 | FECV1 |
| Spike-1 | KP143507 | FIPV1 |
| Spike-1 | KP143508 | FIPV1 |
| Spike-1 | KP143509 | FECV1 |
| Spike-1 | KP143510 | FECV1 |
| Spike-1 | KP143511 | FECV1 |
| Spike-1 | KP143512 | FIPV1 |
| Spike-1 | KX722529 | FECV1 |
| Spike-1 | MH817484 | FECV1 |
| Spike-2 | JN634064 | FECV2 |
| Spike-2 | X06170 | FIPV2 |
| Spike-2 | JQ408981 | FIPV2 |

| | | |
|---|---|---|
| Spike-2 | AB781788 | FECV2 |
| Spike-2 | AB781789 | FECV2 |
| Spike-2 | GQ152141 | FIPV2 |
| Spike-2 | AB907624 | FECV2 |
| Spike-2 | MK987175 | FIPV2 |
