## Supplementary material for "Natural selection differences detected in key protein domains between non-pathogenic and pathogenic Feline Coronavirus phenotypes": SI-TableS2-RecombinantBreakpoints

**Supplementary Table S2.** Inferred GARD recombinant breakpoints.

| Protein | Recombination-free partition (RFP) | Indices (nucleotide) |
|---|---|---|
| Spike-1 | 1 | 1-366 |
| Spike-1 | 2 | 367-729 |
| Spike-1 | 3 | 730-1035 |
| Spike-1 | 4 | 1036-1576 |
| Spike-1 | 5 | 1577-1929 |
| Spike-1 | 6 | 1930-2166 |
| Spike-1 | 7 | 2167-2544 |
| Spike-1 | 8 | 2545-2943 |
| Spike-1 | 9 | 2944-3044 |
| Spike-1 | 10 | 3045-3159 |
| Spike-1 | 11 | 3160-3729 |
| Spike-1 | 12 | 3730-4269 |
| Spike-1 | 13 | 4270-4446 |
| Spike-2 | 1 | 1-138 |
| Spike-2 | 2 | 139-428 |
| Spike-2 | 3 | 429-924 |
| Spike-2 | 4 | 925-1585 |
| Spike-2 | 5 | 1586-1959 |
| Spike-2 | 6 | 1960-3292 |
| Spike-2 | 7 | 3293-4203 |
| Spike-2 | 8 | 4204-4362 |
