## Supplementary figures and images for "Natural selection differences detected in key protein domains between non-pathogenic and pathogenic Feline Coronavirus phenotypes"

### SI-FigureS1-3DstructureORF3c

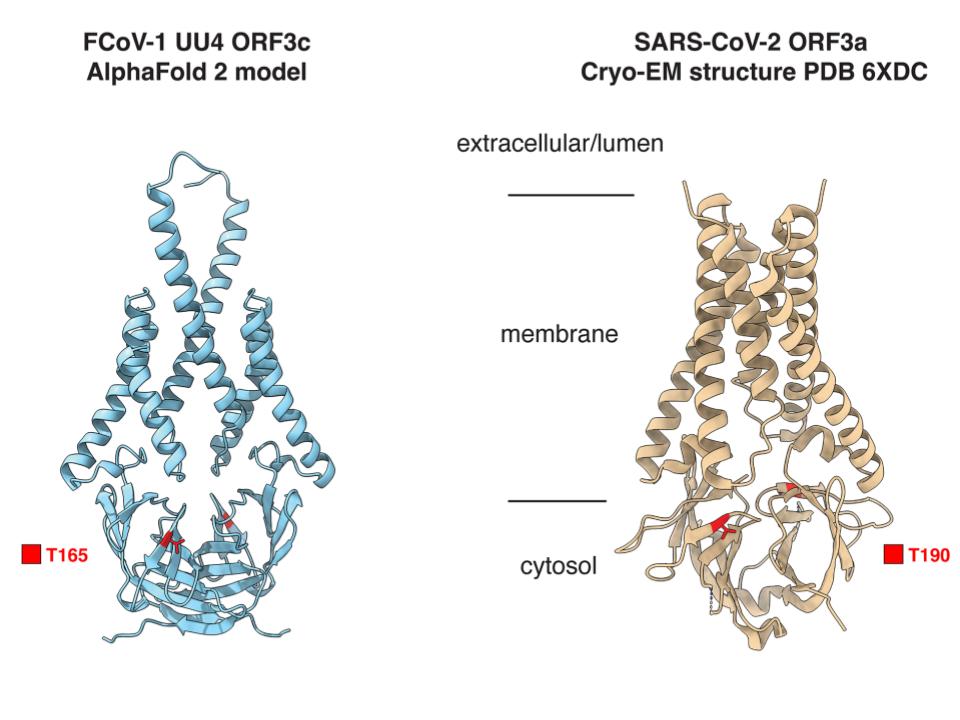

### SI-FigureS2-ProteinAlignment

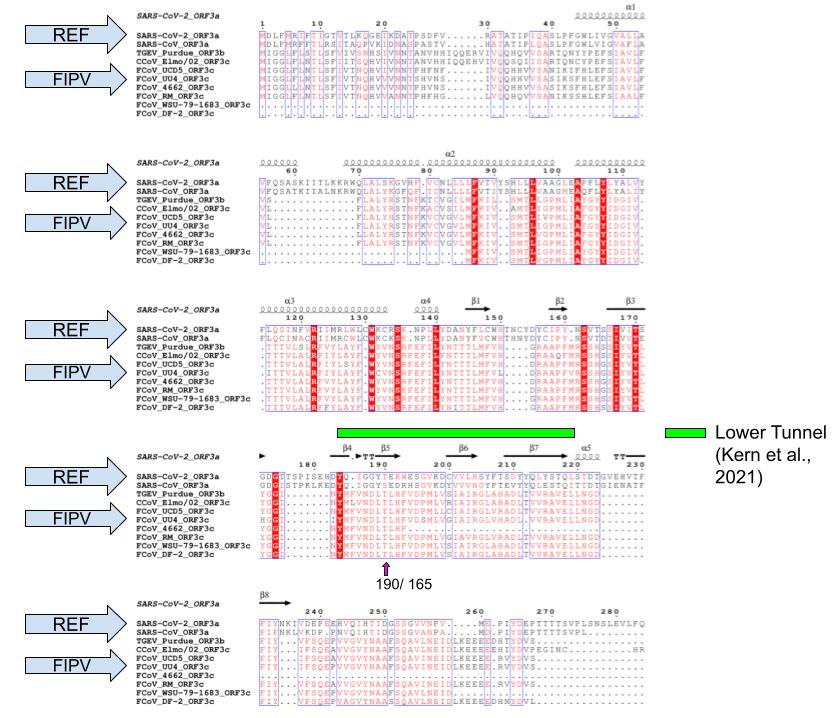
